# Intracellular photonic crystals in photosynthetic sea slugs form via a kidney-mediated biomineralisation pathway

**DOI:** 10.64898/2026.05.07.723475

**Authors:** Samuel Humphrey, Xianglian He, Emeline Raguin, Johannes S. Haataja, Tobias Priemel, Clemens NZ Schmitt, Juliet Brodie, Heather F. Greer, Daniel Wangpraseurt, Lloyd Nelmes, Peter Fratzl, Bruno Jesus, Yu Ogawa, Silvia Vignolini

**Affiliations:** Department of Sustainable and Bio-inspired Materials, Max Planck Institute of Colloids and Interfaces, Am Mühlenberg 1, 14476, Potsdam, Germany; Bio-inspired Photonics Group, Yusuf Hamied Department of Chemistry, University of Cambridge, Lensfield road, CB2 1EW, Cambridge, United Kingdom; Department of Applied Physics, Aalto University School of Science, Aalto, Finland, FI-00076; Natural History Museum, Department of Botany, Cromwell Road, London, UK, SW7 5BD; Scripps Institution of Oceanography, University of California, San Diego, La Jolla, CA 92093, USA; Sea Trust, Ocean Lab, The Parrog, Goodwick SA64 0DE, United Kingdom; Institut Des Substances Et Organismes de La Mer, Département Biologie, Nantes Université, 2 rue de la Houssinière 44322, Nantes, France; Univ. Grenoble Alpes, CNRS, Cermav, 38000 Grenoble, France

## Abstract

Sea slugs in the Sacoglossa superorder are some of the few animals capable of photosynthesising by isolating and maintaining functional chloroplasts within their body^1,2^. While this ability allows some species in this superorder, such as *Elysia viridis*, to appear green, camouflaging themselves within their surroundings^3,4^, this species is marked by extremely bright, coloured regions. Here, we show that these animals produce a yet undiscovered class of photonic structure consisting of intracellular mixed amorphous CaCO_3_ and calcite spherical nanoparticles organised in non-closed-packed face-centred cubic (FCC) lattices and photonic glasses^5^. By mapping the distribution of the cells containing such architectures, we suggest that their colour is linked both to their function and to their biological formation via the animal’s renal system. Using a combination of different optical methods and cryo-electron microscopy, we reveal that the biomineralisation pathway proceeds through stages of calcium ion concentration in the kidney, transport via internal vessels, and precipitation from a dense liquid-like precursor, culminating in the formation of monodisperse nanoparticles, which are the building blocks of these photonic structures.

## Introduction

Sacoglossa are an extraordinary group of sea slugs that engage in kleptoplasty (plastid stealing), a phenomenon in which chloroplasts sequestered from green algae remain partially functional inside the cells of the animals’ sprawling digestive diverticula, providing organic carbon during periods of starvation^1,2^. In *Elysia viridis*^6^, such “stolen” chloroplasts impart a green colouration to their parapodia, which can open and close in response to the light environment^7^, making them resemble leaves^8^. However, as in many other Sacoglossa species^9–12^, these animals are covered by regions of bright iridescent colouration, whose function and composition have so far remained completely unstudied.

Here, we show that these regions of iridescent colouration are cells containing expanded face-centred cubic (FCC) photonic crystal structures composed of calcite-amorphous calcium carbonate (ACC) nanoparticles, which, to the best of our knowledge, have never been reported before and from now on we refer to as ‘calcophores’, in analogy to iridophores^13^. We suggest that they result from calcium ion concentration in the animal’s kidney and subsequent mineralisation from a dense, intracellular, liquid-like precursor. Specifically, by mapping the composition of the precursor and final nanoparticles, we observed that stabilisation of the liquid-like precursor was enabled by the presence of phosphorus and magnesium^14^. We also demonstrate that nanoparticles occupy the entire cell volume and, depending on the filling fraction and degree of packing order, can give rise to different colours and different photonic architectures, i.e. photonic glasses^5^. By mapping the distribution of these photonic structures across the body, we show that their colour is linked both to their function and to their biological formation via the animal’s renal system, composed of the kidney and internal vessels. Specifically, while their position corresponds to the dorsal vessels, their filling fraction is linked to their proximity to the kidney. This gives rise to a high concentration of blue coloured regions above the kidney and more red ones at the edges of the parapodia, with average colour red-shifting along the vessels.

Mineralised photonic structures, such as those in nacre^15,16^ and the blue-rayed limpet^17^, are usually one-dimensional. The only example of an inorganic mineralised 3D photonic crystal architecture is found in Longhorn beetles^18^ (which contain carbonated amorphous calcium phosphate nanoparticles), otherwise only FCC structures composed of organic material have been reported^19,20^. Usually, for the formation of a shell in molluscs, calcium can enter cells via seawater endocytosis^21^ (where cells engulf water directly from their environment), however, it can also be concentrated in the kidney^22–25^. In many Sacoglossa, the kidney is associated with dorsal vessels^26^, which ramify across the parapodial dorsal surface and have both osmoregulatory and excretory function^27–29^. Jensen proposed^4^ that the development of dorsal vessels occurred simultaneously with shell loss. The biomineralisation process in molluscs has been widely studied^21,30–33^, although never in Sacoglossa. So far, it has been reported that ACC typically acts as a precursor for larger mineral structures^16,33–36^, although ACC may be stable in nanoparticle form, such as in cyanobacteria^37,38^, or in bulk (stabilised by phosphate) in crayfish mandibles^39^, lobster cuticles^40^ or as an intermediate calcium storage during the moulting of crayfish in their gastroliths^41^. It is thought to be mediated by a polymer-induced liquid-precursor (PILP)^42–46^ using acidic macromolecules (such as phosphorylated proteins^44,47,48^) or inorganic components such as phosphate or magnesium ions^14,49,50^ to stabilise ACC precursors in a liquid-like phase, although recent studies suggest the PILP phase may be composed of < 50 nm ACC clusters^51^. Once confined^52,53^, the precursor may transition (via transient ACC nanoparticle phases^32,35,36,54,55^) into crystalline calcium carbonate. Biogenic calcium carbonate in marine organisms is found in many polymorphs, including polycrystalline^56^ and amorphous forms, which are determined by the formation process and the presence of additives (organic^36,57,58^ and inorganic^49,59^), thereby disrupting/modifying the crystal lattice formation.

In shells, such as nacre, the highly crystalline mineralised aragonite layers alternate with organic layers, giving rise to a photonic multilayer with optimised mechanical performance^60^. In *E. viridis*, the biogenic calcium carbonate is polycrystalline/amorphous, enabling the formation of spherical nanoparticles and, consequently, a wide range of photonic architectures, from highly ordered photonic crystals to polycrystalline ones and photonic glasses. In this case, polydispersity is hierarchical, present within the nanoparticles, at the nanoparticle assembly level, and in the distribution of the photonic structures. Given the roles structural colour typically serves in other living systems^20,61–65^, we hypothesise that the distinct distribution of these photonic architectures across the body of *E. viridis* may serve a variety of functions, from photoprotection to augmentation of vision and communication.

## Results and discussion

*E. viridis* is covered in bright areas of colour above a background green hue (**Figure 1a**). When viewed at high magnification, these coloured areas are clearly composed of discrete regions (‘calcophores’) lying above the animals’ sprawling green (chloroplast-containing) digestive diverticula (**Figure 1b**). These calcophores are often found in clusters (**Figure 1c**) and exhibit colour across the visible spectrum (**Figure 1d**). Alongside the calcophores, carotenoid pigment granules of a similar size are homogenously distributed across the body^66^. Colour distribution within a calcophore varies: in some cases, it exhibits a distinct texture of different colours, reminiscent of opal gemstones, and in others, it displays a homogeneous colouration throughout (**Figure 1d, Supplementary Figure 1**). Those with an opal-like appearance are iridescent, with colour changing by illumination angle; this effect can also be observed when the animal moves (**Supplementary Videos 1, 2**). Such a varied optical appearance and intense reflectivity (see **Figure 3**) imply that calcophore colour has a structural origin.

**Figure 1.**
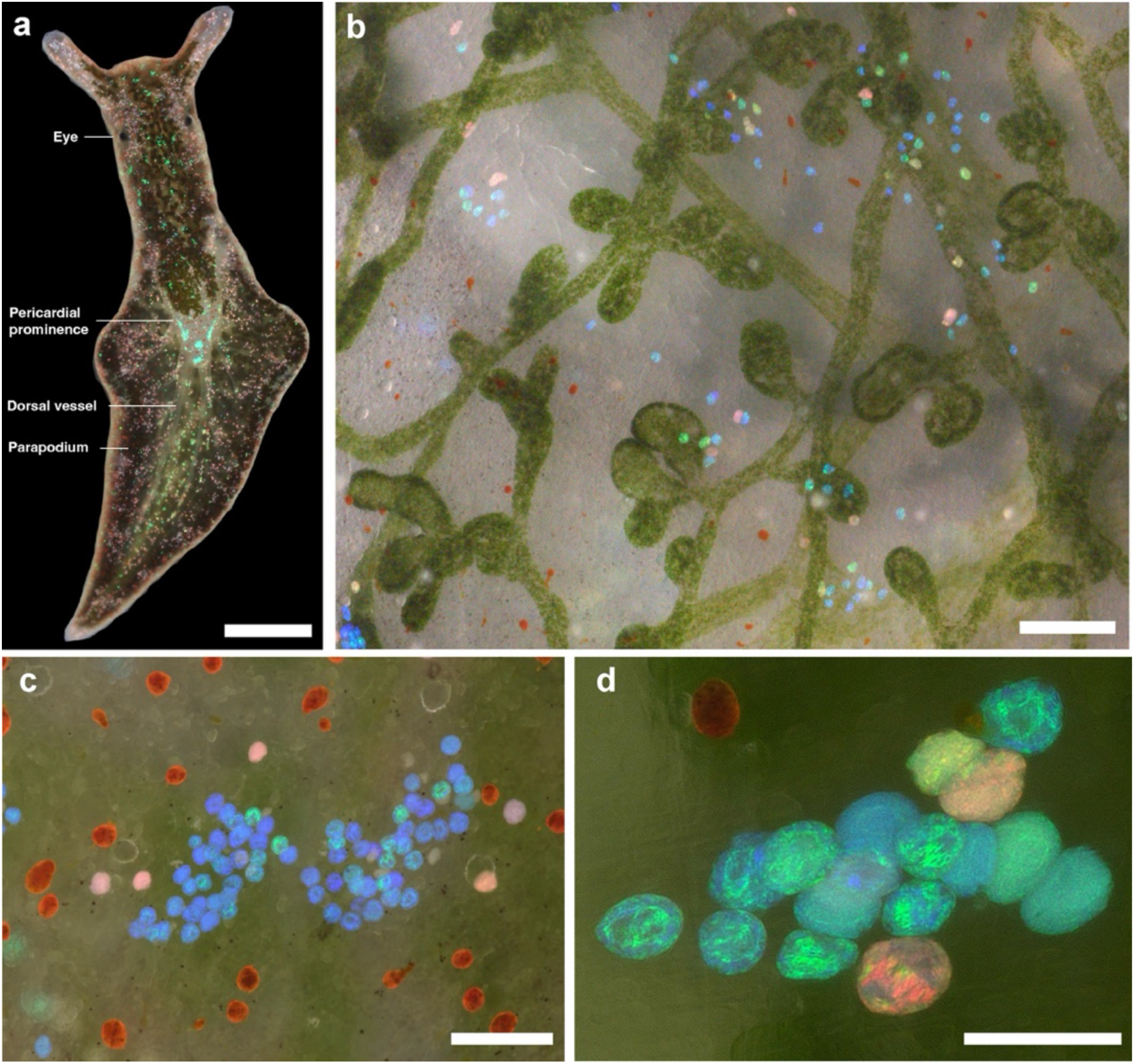
Digital microscopy (Keyence VHX-7100) of *E. viridis* (Sacoglossa, Heterobranchia). a) Whole body, showing bright iridescent calcophores scattered across the body. Red calcophores are concentrated at the edges of the parapodia and around the eye. Blue/green calcophores are concentrated in the centre of the body. The underlying green hue comes from sequestered chloroplasts. Scale bar, 0.1 cm. b) Sprawling digestive diverticula on the dorsal side of an individual, the green hue here comes from sequestered chloroplasts. Calcophores of all colours are scattered above the digestive system. Scale bar, 200 *µ*m. c) Cluster of blue calcophores surrounded by orange carotenoid pigment granules. Scale bar, 100 *µ*m d) Area of calcophores, showing colour variation, and opalescent appearance within one calcophore. Scale bar, 50 *µ*m.

## Ultrastructural analysis

We used electron microscopy to elucidate the nano-architectures responsible for the optical appearance of the calcophores described in **Figure 1**. Using cryogenic focused ion beam scanning electron microscopy (cryo FIB-SEM), we imaged sequential planes of a calcophore from the dorsal parapodial surface and reconstructed a 3D volume from these images (**Figure 2a**). We found an ordered array of nanoparticles (a 3D photonic crystal) with high contrast in the backscattered electron (BSE) image, indicating that the nanoparticles are composed of a heavy-element-rich material. Of the 1791 nanoparticles imaged, we measured an average diameter of 180 nm and a filling fraction of 48.3% (this filling fraction affords space between the particles [a truly close-packed structure has a filling fraction of 74%]). Individual planes showed a high degree of order; for example, **Figure 2a** (inset) shows hexagonal packing in 2D, corresponding to the (111) plane of an FCC structure, with a well-defined interparticle spacing. To elucidate the particle packing in three dimensions, the reconstruction was cropped to an example volume of organised nanoparticles, and 3 sequential (111) planes were shown to stack in the ABC pattern of an FCC structure (**Figure 2a**). Due to the additional space between the particles, we refer to this structure as ‘expanded FCC’. Such an expanded FCC colloid has also been reported to date only in chameleons^20^, though using non-spherical guanine particles. To characterise a larger statistical set of calcophores, we used TEM imaging on cryo-ultrathin sections. Calcophores are found 20 - 100 *µ*m from the surface of the animal and are membrane-bound (**Supplementary Figures 2 and 3a**). Using bright-field TEM, we characterised the morphology of individual nanoparticles, finding that they were not perfectly spherical but rather faceted and rough with a mean nanoparticle diameter of 157 ± 25 nm (*n* = 400), (**Figure 2b, Supplementary Figure 4**). Within an individual calcophore, the polydispersity was low (0.085, measured as the ratio of standard deviation to mean).

**Figure 2.**
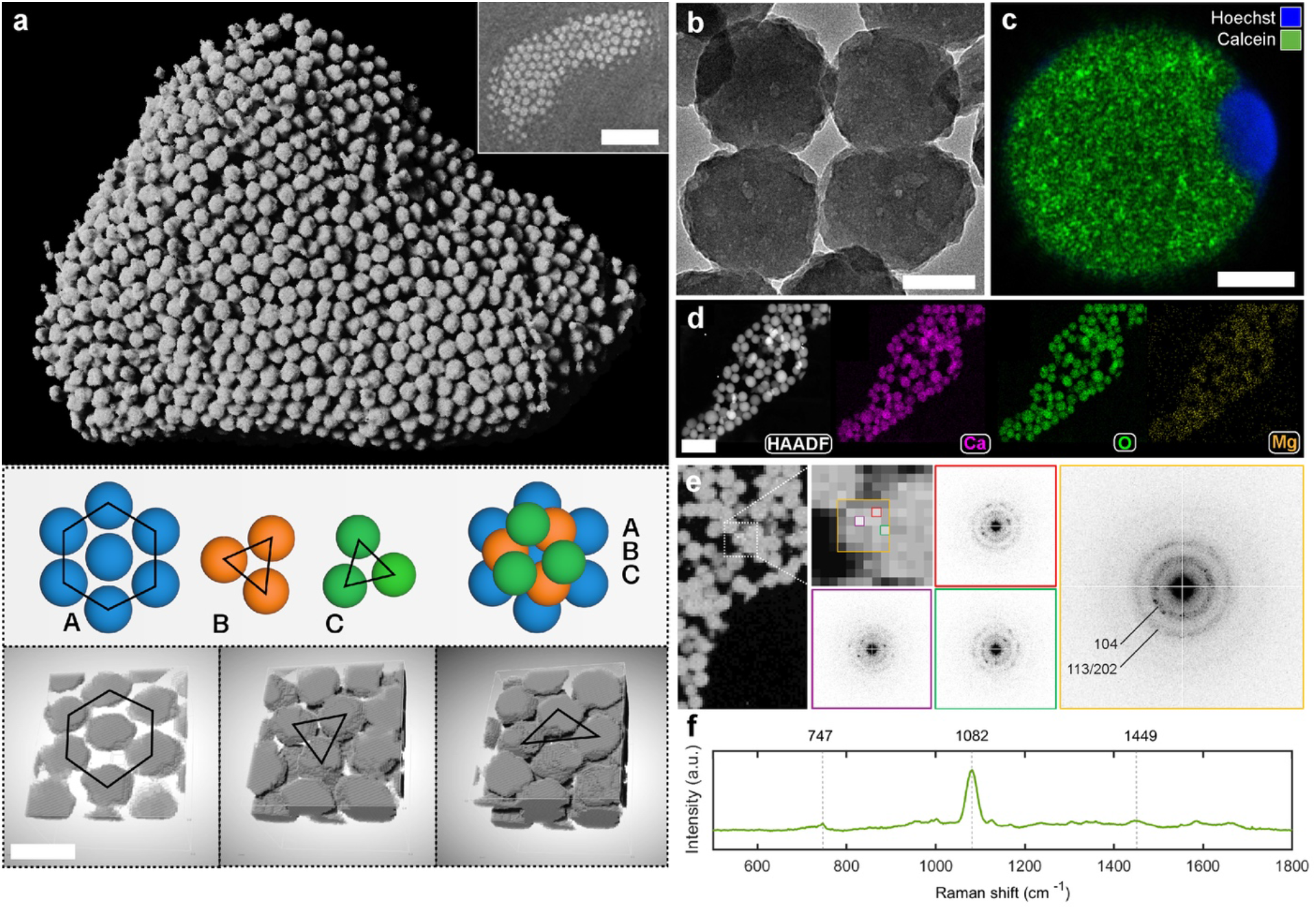
a) 3D cryo FIB-SEM perspective rendering (made using the ORS Dragonfly software) of an *E. viridis* photonic calcophore, showing an array of CaCO_3_ spherical nanoparticles with a high degree of ordering, and space between the particles. Approximate width of region of interest, 7 *µ*m. Inset: Backscattered SEM image of the (111) plane of the expanded FCC structure found in the calcophore, scale bar, 1 *µ*m. Sequential planes of a region of the calcophore in a), showing perfect expanded FCC packing, with an ABC stacking arrangement. Scale bar, 250 nm. b) TEM image showing rough morphology of nanoparticles. Scale bar, 100 nm. c) Confocal microscopy of isolated *E. viridis* calcophore, showing calcein fluorescence from calcium nanoparticles (green) and a nucleus (blue). Scale bar, 5 *µ*m. d) HAADF-STEM image, and Energy dispersive X-ray spectroscopy (EDX) maps of an example *E. viridis* photonic calcophore for the elements: calcium, oxygen and magnesium. Scale bars, 500 nm. e) 4D STEM scanning diffraction analysis of *E. viridis* calcium carbonate nanoparticles. Probe size = 25 nm ø. Diffraction patterns show weak diffraction rings of 104 and 113/202 reflections of calcite I, along with an amorphous halo. f) Raman spectra of photonic calcophores in *E. viridis*. The v_1_ peak (1082 cm^-1^) gives key information on the calcium carbonate polymorph.

To determine the material composition of the calcophores, we used correlated fluorescent staining with the nuclear stain Hoechst 33342 and calcein, along with the surfactant Triton X-100, to make the cell membrane dye-permeable. By imaging stained calcophores isolated from *E. viridis* dorsal surface tissue, we observed that they contain a nucleus and a strong, speckled calcium signal from the remainder of the cell (**Figure 2c**). This matches our TEM results (**Supplementary Figure 5**), where the only two clear components found inside a calcophore were the nanoparticles and an additional nuclear compartment located at the tip of the cell. To verify that the calcium fluorescence originates from the nanoparticles, we acquired STEM-EDS maps from the ultrathin sections. Calcium was the dominating metal with an atomic fraction (*n* = 3) of 17.1%, and oxygen had the highest overall atomic fraction of 79.4%. Carbon was removed from the calculations since the TEM grid had a carbon supporting film. The atomic fraction of oxygen is not entirely representative of the proportion within a nanoparticle, as any organics within the collection area will alter this value. Additionally, the high oxygen content could result from a hydrated mineral phase (typical of ACC^36^). Based on these assumptions, we concluded that the composition of the nanoparticles is hydrated CaCO_3_ with some trace elements such as magnesium (1.4%) and phosphorus (0.4%). Although ACC may be stabilised by phosphate (in lobsters^40^ and crayfish^39,41^), observing stable spherical calcium carbonate nanoparticles is unusual, as they are typically a transitionary state towards the mineralisation of larger structures^33,35,54,67^. To determine the solid-state structure of CaCO_3_ we used high-resolution scanning electron diffraction (4DSTEM). Electron diffraction patterns were obtained by probing multiple areas within a nanoparticle with a 25 nm ø electron probe. The diffraction patterns show broad reflection rings demonstrating that the crystallite size within each nanoparticle is considerably smaller than the probe size (**Figure 2e**). Two diffraction rings observed in the averaged ED pattern have d-spacings of 3.0 and ∼ 2.0 Å corresponding to the calcite I 104 and 113/202 crystal planes^68^. By probing the fine nano and atomic structure of the nanoparticles, we concluded that they are composed of polycrystalline calcite, with a grain size of < 25 nm, and ACC. mThe polycrystalline nature of these nanoparticles explains their rough surface morphology. The magnesium content, along with organic additives and water, may stabilise amorphous CaCO_3_ here, preventing the crystallisation of calcite^45,49,59,69,70^. To further confirm this, we took Raman spectra of calcophores in histological sections. Wehrmeister *et al*. found in their study of biogenic CaCO_3_ that the v_1_ peak lies at 1079 cm^-1^ for ACC, whilst for calcite the peak lies at 1085.5 cm^-1 67^. In the case of *E. viridis* calcophores, the v_1_ peak was broad and lay at the midpoint of the ACC and calcite peaks (1082 cm^-1^ (**Figure 2f**)), confirming the 4D STEM results.

## Optical properties of calcophores

The optical appearance of the calcophores varies not only in colour but also in texture and brightness. This depended on the packing of the nanoparticles. Specifically, we can distinguish two cases: a perfectly periodic packing (photonic crystals) and a correlated one (photonic glasses), in which the average interparticle distance is constant, but the particles are not fixed to well-defined lattice sites. These two categories could be directly distinguished optically as photonic crystals (**Figure 3a**[left]) showed high brightness and texture such as striping, while photonic glasses (**Figure 3a**[right]) had a uniform, dimmer colouration. We attribute the striped texture to twinning or grain boundaries, which create planes of alternating FCC lattice orientations. The result is stripe-like coloured motifs^71^. However, as more clearly shown in **Supplementary Figure 1**, within a single calcophore, we can recognise features of both categories. The fact that the degree of structural ordering not only varies within a single calcophore, but also between them, can be further confirmed by TEM imaging. Many calcophores show amorphous packing of the nanoparticles (**Supplementary Figure 3b**), polycrystalline packing (**Supplementary Figure 3c**), or mixed amorphous and crystalline packing (**Supplementary Figure 3d**). The ordered domains may extend up to 6 *µ*m, typically from the edge of the calcophore towards the centre.

**Figure 3.**
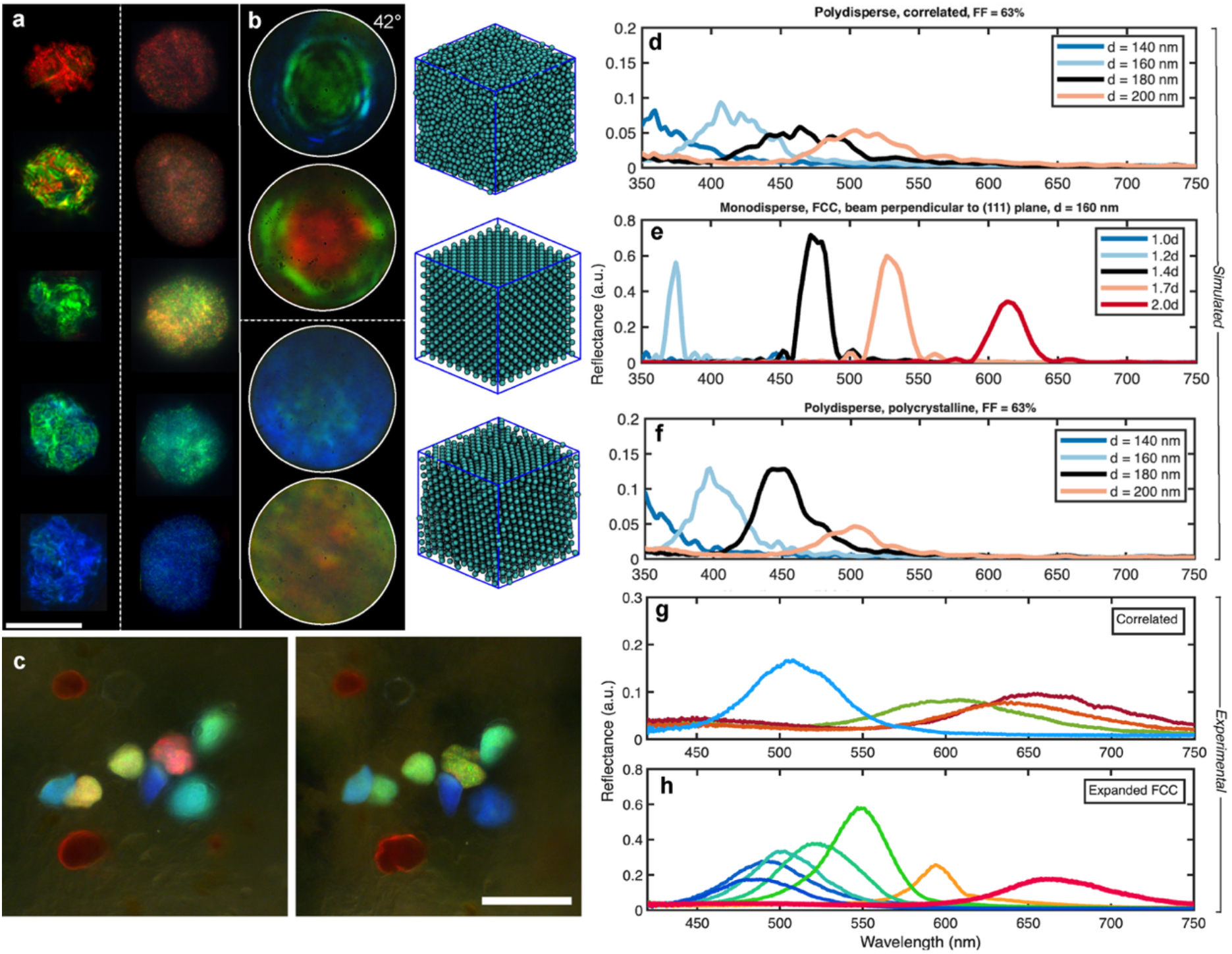
a) Bright-field upright optical microscope (Zeiss axioscope, 63X WI objective) of calcophores on the dorsal surface of *E. viridis*, showing both homogenous ‘glassy’ colour (right), and striped, opalescent ‘crystalline’ optical texture (left). Exposure time for ‘glassy’ images is 2.5X that of ‘crystalline’ ones. Scale bar, 20 *µ*m. b) Reciprocal space (K-space) images for calcophores showing both ‘glassy’ (bottom) and ‘crystalline’ (top) type optical appearance corresponding to real space images in **a**, glassy K-space images show homogenous colouring, with a very slight blue shift at high angles. Crystalline K-space images show a strong blue shift at higher scattering angles, and a highly textured, hexagonal colour distribution corresponding to the angular response of the photonic crystals. The edge of the circle in the K-space images corresponds to NA = 0.9 (scattering at 42^°^). Images taken with Zeiss axioscope, 63X WI objective. c) Dark-field upright optical microscope images of a cluster of calcophores before and after 20 minutes exposure to air. Dark orange carotenoid volumes (top and bottom left) can be used as markers to correlate the calcophores in the two images. Of the 7 calcophores in the cluster 4 of them show a blue-shift. Scale bar, 50 *µ*m. FDTD simulations of 5 *µ*m diameter spherical photonic structures found in *E. viridis*. A gaussian beam of radius 1.25 *µ*m was used for all simulations. The particles have been simplified to perfect spheres. d) Simulated spectra for a correlated structure (photonic glass), with particle sizes varying from 140 - 200 nm. e) Simulated spectra for expanded FCC photonic crystal with particle-to-particle distances varying from 1 d to 2 d (where d is the diameter of one particle, in this case 160 ± 5 nm). The (111) plane is perpendicular to the detector for simulations. f) Simulated spectra for close packed spheres with polycrystallinity i.e. the simulated structure is composed of multiple crystallographic volumes for d = 140 - 200 nm. g) Experimental bright field spectra (Zeiss axioscope, 63X WI objective, 50 *µ*m fibre) spectra of *E. viridis* photonic calcophores, showing photonic glass-like response. Spectra are normalised to a silver mirror. h) Experimental bright field spectra (Zeiss axioscope, 63X WI objective, 50 *µ*m fibre) spectra of *E. viridis* photonic calcophores, showing photonic crystal-like response. Spectra are normalised to a silver mirror.

**Figure 3b** shows reciprocal space (K-space) images of crystalline and glassy calcophores, respectively. In these images, the outer ring of the circle is defined by the numerical aperture of the objective lens, NA = 0.9, and the centre corresponds to normal incidence, NA = 0 (angle increases linearly with arcsin of the distance from the centre of the image, from 0^°^ to 42^°^). As expected, calcophores with a photonic glass-like response showed homogenous colour across the whole K-space image, with a slight blue-shift away from the centre. For photonic crystal-like calcophores, the K-space images showed hexagonal motifs with strong colour variation and blue shift at increasing angles due to the strong angular dependence of photonic crystals.

In addition to the particle arrangement, the filling fraction or interparticle spacing influences photonic properties of the calcophores. To probe this effect, we imaged a cluster of calcophores before and after 20 minutes of dehydration by exposure to air. **Figure 3c** shows that the majority of calcophores blue shifted due to structural contraction, this effect was also clear in isolated calcophores (**Supplementary Figure 6**).

To match the optical properties of the photonic structures described here to those observed by electron microscopy and to demonstrate an expanded-FCC lattice that captures the measured optical response, we used finite-difference time-domain (FDTD) calculations on structures generated by molecular dynamics^72^ (**Figure 3d-f**). We did not attempt to match simulated spectra 1-to-1 with experimental spectra, but rather compared the salient spectral features, such as the intensity and width of the reflection peaks, across different degrees of order. By comparing the numerical calculations reported in **Figure 3d-f** with the experimentally measured reflections from the individual calcophores in **Figure 3g, h**, we concluded that: depending on the calcophore, and region within a calcophore, only the simulated responses of an expanded FCC, polycrystalline expanded FCC, or correlated structure are well capable of capturing those salient spectral features observed experimentally, and that a decrease in filling fraction causes a red shift in the optical response, see **Supplementary section 3** for expanded figures/discussion. For the FDTD calculations, we used a refractive index of 1.58 [average value of (i) ACC and (ii) birefringent calcite^73,74^], giving contrast to the matrix material, assumed to have a refractive index of 1.34 (typical of cytosol in marine organisms).

## Interpreting calcophore distribution

To determine whether the colour distribution of calcophores might have a biological role, we analysed their distribution across the body by colour. Visually, the dorsal distribution of calcophores differs from the ventral one (**Figure 4a** and **Supplementary Section 4**). More precisely, we can identify 6 distinct areas with different colour patterning (**Figures 4b, c**).

**Figure 4.**
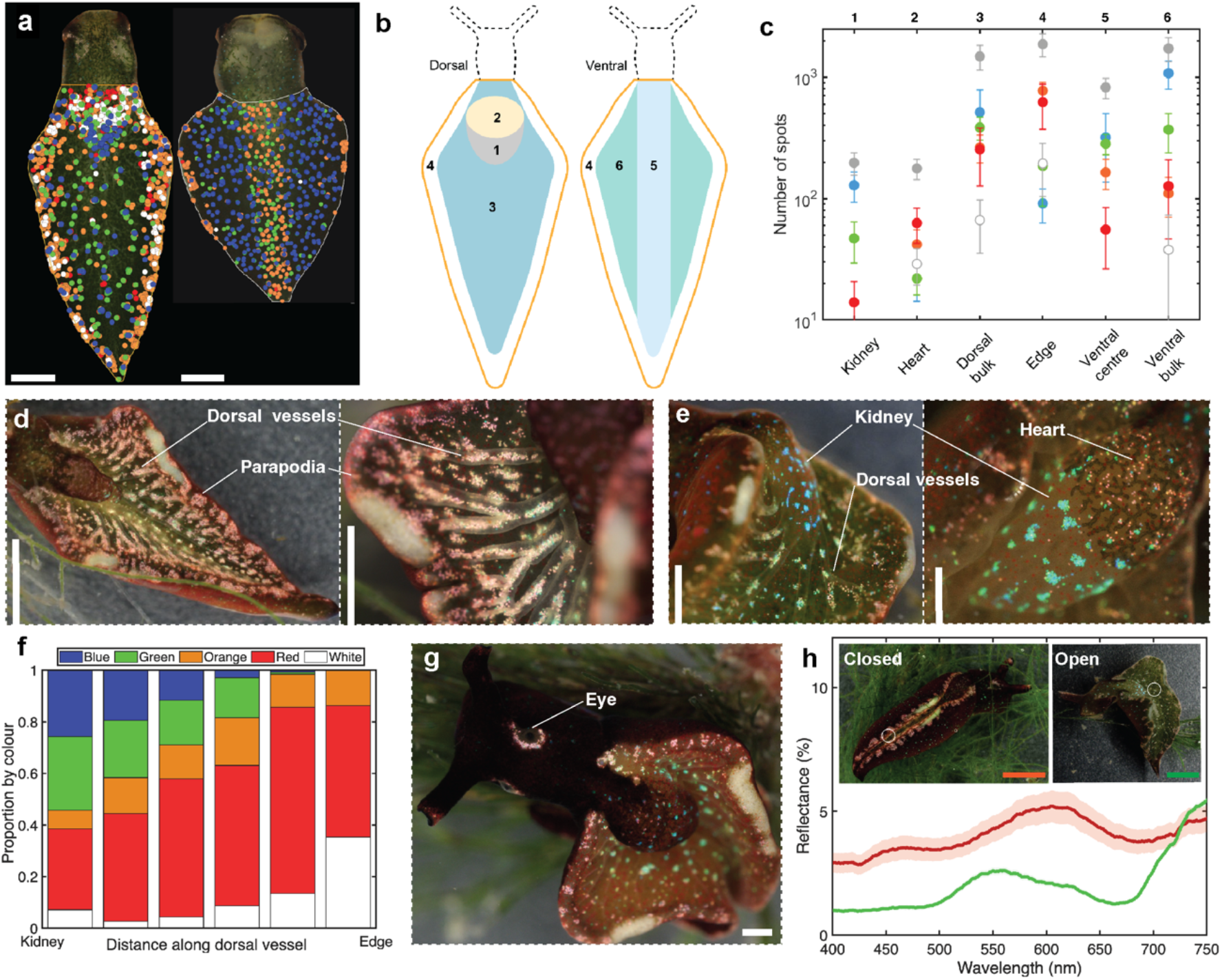
a) Example images of the dorsal and ventral sides of an *Elysia viridis* individual. Orange, red, white, green and blue calcophores are mapped as coloured dots using the QGIS mapping software. Variation in pattern between colours is visible, such as the frequency of orange/red calcophores at the parapodial edge, or blue calcophores on the posterior side of the pericardial prominence. The head of the animal was mapped separately. Scale bars, 0.5 mm. b) 6 distinct regions defined by body mapping, these are: kidney (posterior pericardial prominence, area 1), heart (anterior pericardial prominence, area 2), dorsal bulk (area 3), edge (area 4), ventral centre (area 5), ventral bulk (area 6) c) Colour mapping for the 6 regions defined in b) for the colours blue, green, orange, red, white and the total (marked in grey). Values are averaged over 6 individuals, and error bars represent standard deviation. Y axis has a log scale. d) Digital microscope (Keyence VHX-7100) images of the dorsal surface of an individual, showing the clear correlation of dorsal vessels to calcophores, and high density of red and white calcophores at the edge of the parapodia. Scale bars, 2 mm, 0.5 mm. e) Digital microscope images of the dorsal pericardial prominence (heart & kidney) of an individual, showing the high density of blue calcophore clusters above the kidney region, discrete red calcophores above the heart and red shift of calcophores along dorsal vessels away from the kidney towards the parapodial edge. Scale bars, 0.6 mm, 0.5 mm. f) Histogram representing the proportion of each colour of calcophore along a dorsal vessel from the kidney to the parapodial edge (excluding the kidney and edge themselves). There is a clear red shift in calcophore colour along the vessels. g) Example image of an individual with a clear ‘ring’ of red calcophores around the eye. Scale bar, 1 mm. h) Double-ended probe measurements of an *Elysia viridis* individual with parapodia open and closed, showing the change in spectral response due to the photonic calcophores on the edge of the parapodia, which we suggest have a photoprotective role. Spectra are normalised to a silver mirror. The inset shows Keyence microscope images of *Elysia viridis* individuals in the closed (with a clearly visible strip of photonic calcophores) and open morphologies. Scale bars, 2 mm.

We observed a high concentration of clustered blue calcophores on the posterior side of the pericardial prominence (kidney, area 1), the dominance of individual red/white/orange calcophores on the anterior side of the pericardial prominence (heart, area 2), the high density of red/white/orange calcophores on the edge of the parapodia typically in clusters which bridge the dorsal and ventral sides (area 4) (**Figure 4b**). These patterns are shown in the example images **Figure 4d, e**. The other patterns are less clear without point maps (as highlighted in **Figure 4a**); a high density of clustered blue/green calcophores in the bulk of the surface area on both the dorsal and ventral sides (areas 3 and 6), and a strip of calcophores along the ventral side, which was red-shifted from the calcophores in the parapodia (area 5).

In all individuals, we observed that the colour of calcophores near the kidney region was blue-shifted relative to those at the parapodial edges. In individuals with an especially large number of calcophores, there was a clear correlation between the position of the calcophores on the dorsal side and the dorsal vessels, which protrude from this surface (**Figure 4d, e**). These vessels originate at the pericardial prominence and ramify across the surface. This has also been reported in other species of Sacoglossa^9^. In individuals where this effect is clear, following the vessel away from the kidney to the edge of the animal, the colour of calcophores red-shifts (**Figure 4e**). For individuals where the density of calcophores is not high enough to trace a continuous line under the vessels, the colour shift is still clear (**Figure 4g**). Blue calcophores have a higher density of nanoparticles than red ones, thus the mineralisation process may be linked to their colour distribution. The high density of blue calcophores at the kidney suggests a high degree of mineralisation in this region, resulting in calcophores with a high filling fraction and, therefore, photonic crystals that reflect blue light. The connection of dorsal vessels to the kidney region (**Supplementary Figure 13**) and their correlation to the calcophore positions suggest that they may act as transport vessels for mineralisation precursors. If this is the case, we hypothesise that calcophores in proximity to the kidney have access to higher concentrations of mineral precursors with respect to those at the end of these vessels, resulting in a lower rate of mineralisation with decreasing proximity to the kidney and therefore a lower filling fraction of nanoparticles (and red-shifted optical response). To support the hypothesis that colour is linked to the rate of biomineralisation (which we believe is related to the internal concentration of calcium ions), we used body regeneration^75^ as a proxy for initial body growth and calcophore formation, see **Supplementary Figure 14**.

Whilst we suggest the colour and position of calcophores is intimately linked to their biogenesis, we also believe that their distribution is functional. Heads were mapped separately from the body and images were taken whilst the slugs were free roaming (**Supplementary Figure 15**). In every individual, there is a ‘halo’ of calcophores around the eye, which are exclusively red/white. Because this is consistent across all individuals studied, we suggest it may be functional, augmenting vision by increasing the amount of light reaching the eye via scattering or by increasing the conspicuousness of the animal’s head for orientation during mating. The locations of bright regions of calcophores, such as above the kidney, or red calcophores on the perimeter of the parapodia, could also act to increase the conspicuousness of these regions to other individuals, helping with orientation during mating.

The high density of calcophores at the parapodial edges of *E. viridis* is consistent across every individual. Kleptoplasts inside the cells of *E. viridis* are highly vulnerable to light damage, and it has been reported in multiple species, including *E. viridis*, that animals will close their parapodia in response to high light^7,76^. Closing the parapodia reduces the animal’s overall surface area, reducing light exposure to chloroplasts, but it also directs the clusters of calcophores on the parapodial edges upwards. If this region of the animal is approximately 1 cm long, then the result is roughly 1 mm^2^ of area covered by reflective calcophores, see **Supplementary Section 4** text. This could be a key factor, amongst others^77,78^, which allows this species to extend kleptoplast lifetime. By closing the parapodia the animal can selectively limit light to the chloroplasts, under low light conditions maximising solar irradiation is key for the chloroplasts. To quantify this effect, we took large area average reflectance spectra of the animal in the open and closed parapodial positions (**Figure 4h inset**). **Figure 4h** shows spectra taken of an individual with parapodia open and closed, showing that the reflective peak in the red region in the closed position effectively increases back-scattering, approximately doubling the total amount of reflected light. We also note an additional structurally coloured reproductive organ, elaborated on in **Supplementary section 5**.

## Calcophore formation

To understand the process of calcophore formation, we imaged newly formed parapodial tissue and correlated images of the same area using different optical techniques. Specifically, we used images obtained in reflection mode (R) and transmission mode (T) with a digital microscope, and holotomographic images (HT), see methods. While we cannot directly track the live development of a calcophore, **Figure 5a-e** reports what we believe represent different stages of calcophore development, after having observed consistent features in several individuals of different ages. A fully developed calcophore (**Figure 5a**), which was brightly coloured in R, appears as a textured isolated region in HT. In contrast, a more homogeneous texture and a decrease in reflected colour strength were observed as we go from **Figure 5b** to **Figure 5d** (indicative of stages prior to the formation of a photonic structure). In these cells inner *and* outer membranes were visible and the cells were not isolated but connected to a calcophore ‘conduit’, which we suggest is responsible for the transport of mineralisation precursors into the cell and may be connected to the dorsal vessels ramifying from the kidney. To confirm the hypothesised calcium transport link between the kidney, dorsal vessels, conduits and calcophores, we recorded the calcium distribution across the body by staining live individuals and histological sections with calcein (**Figure 5f, Supplementary Figure 21** respectively). Specifically, we observed (using confocal microscopy to map the fluorescent signal of calcium): (i) a high fluorescence signal in the kidney and dorsal vessels, and (ii), when viewed under high magnification, regions that resemble the shape and size of calcophores and their conduits observed in HT (containing a discrete volume of calcium) in the parapodia (**Figure 5f inset**), see also **Supplementary Figures 21, 22**. This whole-body calcium mapping supports our hypothesis that calcium is concentrated in the kidney and subsequently transported via vessels to the calcophores, and explains the positional correlation of the calcophores with respect to both the kidney and dorsal vessels (**Figure 4**). Calcein is cell membrane impermeable and therefore is likely to have reached the volume in **Figure 5f** (inset) via the conduit. Therefore, fully formed calcophores are not visible with in vivo calcein staining and must be isolated and permeabilised with a surfactant to be imaged.

**Figure 5.**
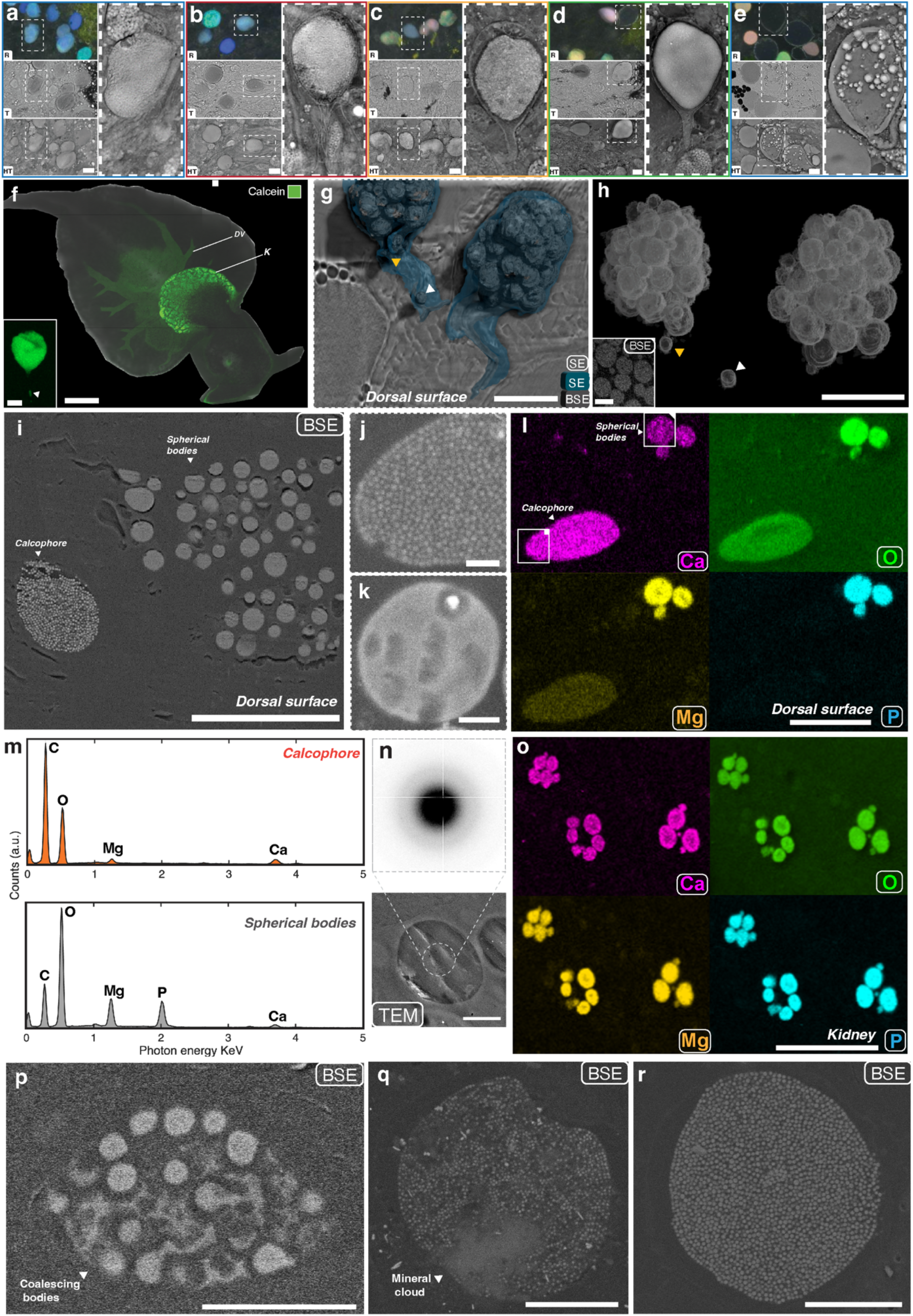
Correlated optical microscopy of newly grown parapodial tissue: Top (**R**) = Reflectance (Keyence Full-Ring mode), Middle (**T**) = Transmission (Tomocube HT-X1™), Bottom (**HT**) = Holotomography. Scale bars, 15 *µ*m. a) Fully formed blue calcophore. Calcophore appears blue in reflectance mode, dark in transmission mode (due to reflected light decreasing signal). Holotomography shows nanoparticulate texture throughout, nucleus at the top of the cell, an outer membrane, and no other cellular components. b) Calcophore with blue-white appearance in reflectance mode, dark appearance in transmission mode. Holotomography shows nanoparticulate texture (throughout majority of cell, there are sparse areas in the bottom of the cell), nucleus at the top of the cell, an outer membrane, inner membrane and vessel like appendage at the bottom of the cell. c) Calcophore with white appearance in reflectance mode, dark appearance in transmission mode. Holotomography shows nanoparticulate texture (contained within an inner membrane), nucleus at the top of the cell, an outer membrane/cell wall, and vessel like appendage at the bottom of the cell. d) Cell (possible calcophore precursor) which appears transparent in both reflectance and transmission modes (due to lack of photonic response because of homogenous internal structure [no nanoparticles]) e) Cell (possible calcophore precursor) which appears transparent in both reflectance and transmission modes, containing discrete spherical bodies contained within an inner membrane, and inside the attached vessel. f) Tiled confocal image (Zeiss LSM 880) of a sedated, live *E. viridis* individual stained with caclein (green), showing high degree of calcein fluorescence in the kidney region, and in the dorsal vessels ramifying from the kidney. Scale bar, 0.2 mm. K = kidney, DV = dorsal vessel. Inset shows high magnification confocal (Zeiss LSM 980) Z projection of an isolated volume within the parapodia of the same individual. The morphology of this volume matches well with the holotomographic images in **b**-**d**. Additionally a calcium containing vessel is visible extending from the bottom of the volume. Scale bar, 5 *µ*dm. g) Combined 3D reconstruction of a cluster of high BSE signal spherical vesicles (segmented from BSE images [grey]), and the matrix material surrounding them which is directly connected to a conduit (segmented from inLens SE images [blue]). High BSE spherical bodies are visible within the left vessel (see white/yellow arrows). A single SE image is included for context. The volume imaged here is at the dorsal surface of an adult *E. viridis*. Scale bar, 5 *µ*m. h) 3D reconstruction of only the BSE signal from the spherical bodies, demonstrating their sphericity and clustering, as well as the isolated bodies, which may be transported towards the cluster. Scale bar, 5 *µ*m. Inset shows a single BSE image from which the reconstruction was made. Scale bar, 1 *µ*m. i) Block face SEM in BSE mode of freeze substituted and embedded dorsal surface tissue, showing a calcophore (left) in close proximity to a cluster of high BSE bodies. Scale bar, 10 *µ*m. j) Cropped block face SEM of part of a calcophore, from a region of interest (ROI) containing both a calcophore and spherical bodies. Scale bar, 1 *µ*m. k) Cropped block face SEM of a single spherical body taken from the same ROI as j. Scale bar, 1 *µ*m. l) EDS maps for the elements Ca, O, Mg, and P for the ROI containing both a calcophore and cluster of spherical bodies. Ca, O and Mg are present for both the calcophore and the bodies, whilst P is only present in the bodies, implying its involvement in preventing precipitation of nanoparticles. Scale bar, 10 *µ*m. m) EDS spectra for the calcophore and spherical body in I, showing similar spectral peaks for all elements aside from P, which is only present in the spherical bodies. n) TEM image and diffraction pattern of a precursor body showing an amorphous halo with no distinct diffraction spots, implying an amorphous/liquid-like atomic structure. Scale bar, 1 *µ*m. ø = 800 nm. o) EDS maps for the elements Ca, O, Mg, and P for a ROI inside the kidney of an adult *E. viridis*. Clusters of spherical bodies are visible with elemental maps corresponding to the bodies found in the parapodia. Scale bar, 25 *µ*m. p) Block face SEM of freeze substituted and embedded dorsal surface tissue, showing a cluster of high BSE spherical bodies, some of which have ruptured, and are spreading out with fluid-like patterns. Scale bar, 2.5 *µ*m. q) Block face SEM of freeze substituted and embedded dorsal surface tissue, showing an array of nanoparticles, smaller than typical for a fully formed calcophore, and a ‘cloudy’ area of high BSE signal material, containing no nanoparticles, possibly because it is from this material that nanoparticles precipitate. Scale bar, 5 *µ*m. r) Block face SEM of freeze substituted and embedded dorsal surface tissue, showing a fully formed calcophore. Nanoparticles are notably larger than in q. Scale bar, 5 *µ*m.

Finally, we suggest that **Figure 5e** shows a calcophore cell with a conduit resembling **Figures 5b-d**, however, here the inner membrane contains micron-sized (2.1 ± 0.9 *µ*m diameter, *n* = 42) ‘spherical bodies’ (an order of magnitude larger than the spherical nanoparticles observed in a fully formed calcophore [**Figure 5a**]), which may be loaded inside the cell via the conduit. We propose that these spherical bodies dilute and coalesce in response to chemical changes in the intracellular environment, such as pH change or rehydration^54,79,80^ (resulting in a morphology resembling **5d**) prior to the onset of mineralisation (resulting in morphologies resembling **Figure 5b, c**).

To understand the biomineralisation mechanism in calcophore development, we imaged dorsal parapodial tissue both in its native state using cryo-FIB SEM on high-pressure frozen tissue (**Figure 5g, h**) and (along with kidney tissue) SEM on freeze-substituted tissue. We located a volume in FIB SEM containing high BSE signal spherical bodies (1.1 ± 0.5 *µ*m diameter, *n* = 42). In the image reconstruction obtained from the SEM images, we found that these bodies are bound within a membrane, which is directly connected to a conduit (see **Supplementary Figure 23** for 2D images). Inside the conduits, additional spherical bodies were visible, suggesting that they may be transported into the calcophore cells via the conduits prior to nanoparticle formation.

Block face SEM revealed similar clusters of high BSE signal spherical bodies, adjacent to fully formed calcophores, close to the parapodial surface (**Figure 5i**) with a mean diameter 1.2 ± 0.4 *µ*m (*n* = 42), these clusters contain tens of spherical bodies (**Supplementary Figure 24**). Using EDS mapping, we compared the elemental composition of a fully formed calcophore and a spherical body in the same region of observation in the SEM field of view. While the nanoparticles and spherical bodies both contain Ca, O and Mg (with a higher signal in the bodies), P is only meaningfully present in the spherical bodies (**Figure 5l, m**). We also observed spherical bodies in the kidney tissue (here the clusters contain just a few (**Figure 5o**)). EDS mapping revealed near-identical composition in these two regions (*Parapodia* (at%), O:48, P:17, Ca:6, Mg:12, *Kidney* (at%), O:49, P:19, Ca:5, Mg:13, with results averaged over 3 discrete clusters for each region). The high P^81^ (which may be associated with proteins, or small molecules^14,47,48,50^) and Mg^14,49,59^ content stabilise this liquid-like amorphous state, as demonstrated in lab-based PILP systems^47,82–84^, but also without additives^85,86^. The sphericity of these micron-scale bodies is consistent with hydrated amorphous precursor phases, which lack crystallographic anisotropy and behave in a liquid-like manner where minimising interfacial free-energy promotes spherical morphology^87,88^. This is further confirmed by electron diffraction of the spherical bodies (**Figure 5n**), which revealed an amorphous halo, with no distinct diffraction spots. We found no evidence through EM of a membrane bounding individual spherical bodies and therefore suggest that they may be liquid-like condensates, rather than membrane-bound vesicles. This would allow for their dissolution in response to changes in the local chemical environment within the calcophore membrane.

We therefore hypothesise that, prior to mineralization, these bodies diffuse and coalesce (as observed in **Figure 5p**, which suggests fluid-like behaviour of coalescing bodies in the BSE mode) in response to changes in the local environment such as pH shifts or rehydration, releasing their contents. The high local concentration of Ca, Mg, P, and macromolecules could demix, exposing Ca^2^+ ions to chemical triggers such as carbonate ions, initiating mineralisation into solid nanoparticles (via liquid-liquid phase separation [LLPS]^82,83,89^). **Figure 5q** shows a calcophore containing smaller than average (e.g. vs **Figure 5r**) nanoparticles (< 125 nm) as well as a high BSE signal ‘cloudy’ area containing no nanoparticles. This may represent the mineral-rich liquid-like phase prior to the onset of LLPS. The isolation of this phase within the inner calcophore membrane allows for close control over the local chemistry, which is determinate in the final CaCO_3_ phase, permitting precise control of nanoparticle morphology and stability^37,85^.

In summary, we propose that specialised kidney cells accumulate intracellular condensates/bodies loaded with high concentrations of Ca^2+^, phosphate, Mg^2+^, and/or phosphorylated proteins or other macromolecules. These mineral-loaded bodies are transported via ramifying vessels to the calcophore cells, which induce dissolution by altering the local chemical environment, destabilising the liquid-like mineral precursor phase, and promoting LLPS-mediated mineralisation to form organised CaCO_3_ nanoparticles. These nanoparticles are the building blocks of the fully formed photonic architectures we observe. While more research is required to confirm the biological significance of these structures, we believe that mineralisation pathways remnant after shell loss may have been co-opted to generate structurally coloured and optically functional mineralised structures in Sacoglossa.

## Methods

The detailed methods section has been included in the **Supplementary Information Section 7**.

## Supporting information

Supplementary figures 1-31 and text sections 1-7

## Acknowledgements and funding

This work was supported by the ERC BiTe ERC-2020-CoS-101001637 to SV, SH, the EU Horizon 2020 program (H2020-MSCA-ITN-2019) grant N 860125 “BEEP” for SV and BJ, JB, the Marie Skłodowska-Curie grant agreement No.101155041 to SV and TP, the Marie Skłodowska-Curie grant agreement No.893136 to SV and J.S.H., the Emil Aaltonen Foundation and Academy of Finland grant (no. 347789) to J.S.H. SV acknowledges the Max Planck Society for the generous financial support. LN acknowledges Sea Trust, Wales. The authors would like to acknowledge Junwei Wang for analysis of colloidal packing, Gea van de Kerkhof for project initiation, Maria Murace for optical microscopy assistance, Kathe Jensen for advice on physiology, Michael Kühl and Swathi Murthy for suggesting the use of Tomocube, Alon Gorodetsky for critical feedback on the results and for suggesting performing the staining of the slugs. SH acknowledges Edward Wills for training on histological staining. We finally acknowledge Heather Greer, Darran Clements and Louis Elfari for access to the EM facilities at the University of Cambridge and EPSRC for funding the TEM (EP/P030467/1). YO acknowledges Agence Nationale de la Recherche (ANR grant number: ANR-21-CE29-0016-1) for the financial support and the NanoBio-ICMG platform (FR 2607) for granting access to the electron microscopy facility. The work has benefited from characterisation equipment of the Grenoble INP-CMTC platform supported by the Centre of Excellence of Multifunctional Architectured Materials ‘CEMAM’ n°ANR-10-LABX-44-01 funded by the Investments for the Future program.

