## Supplementary figures 1-31 and text sections 1-7 for "Intracellular photonic crystals in photosynthetic sea slugs form via a kidney-mediated biomineralisation pathway"

### 1. Digital microscopy

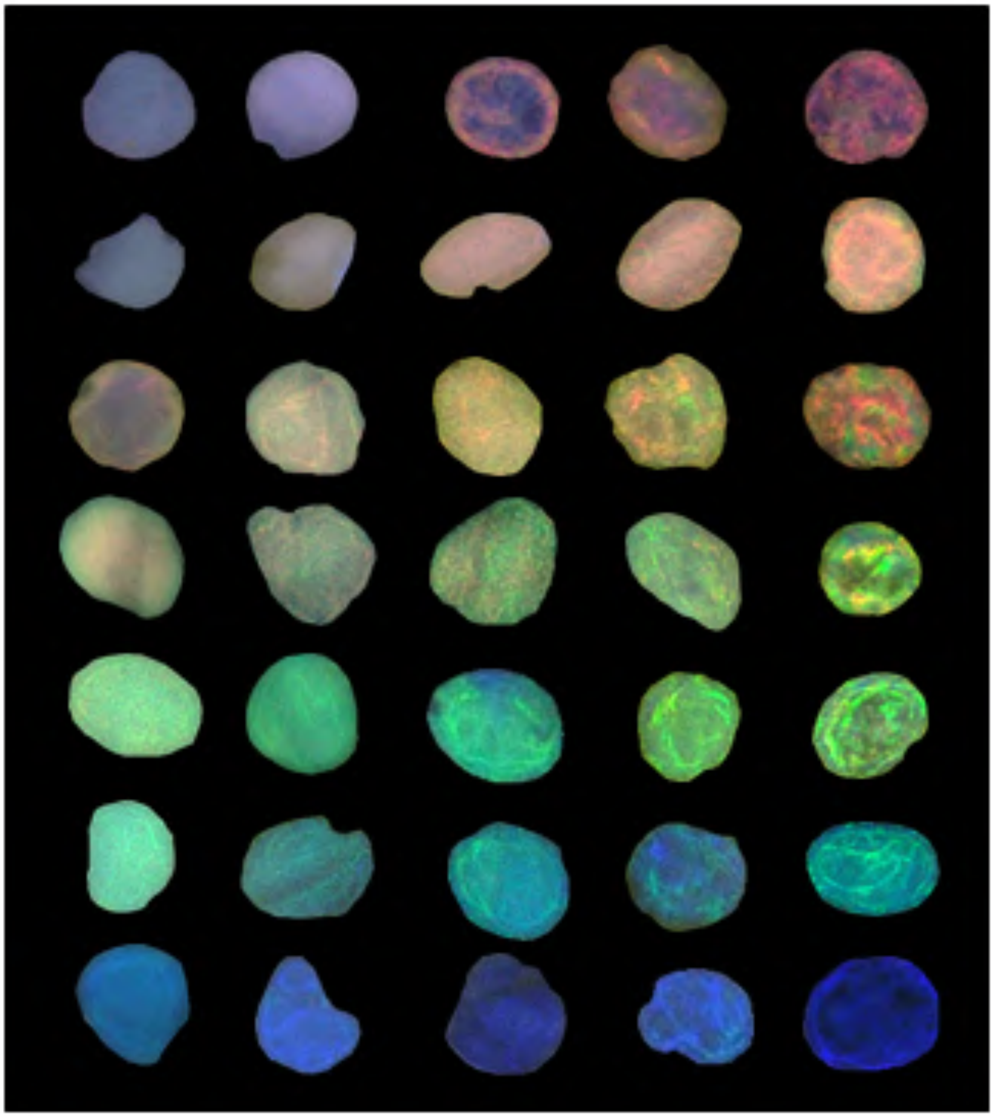

**Supplementary Figure 1**

Digital microscope images of in vivo calcophore of *Elysia viridis* parapodial tissue. Cells show a range of morphologies in two dimensions and a variety of optical textures.

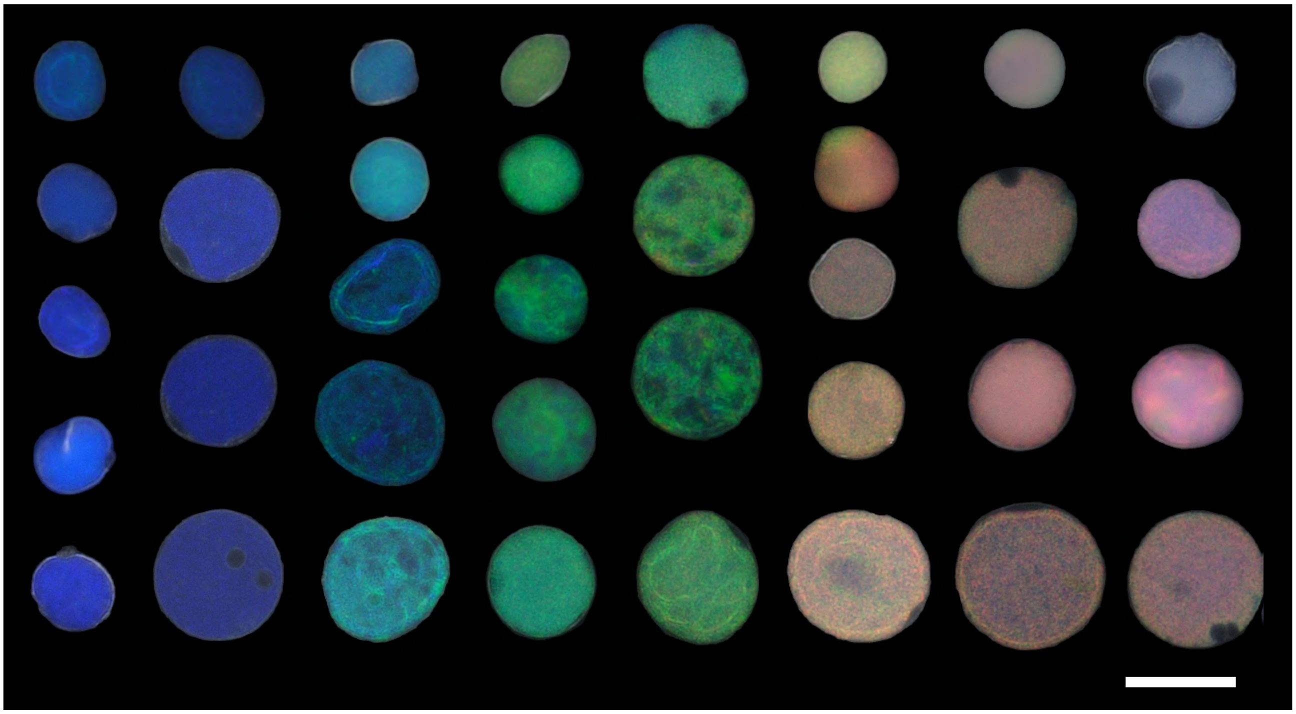

**Supplementary Figure 2**

Digital microscope images of isolated calcophores of *Elysia viridis* parapodial tissue. Cells show circular shape in two dimensions. Scale bar, 25 µm.

Viewed in 2 dimensions, in vivo calcophores are often oblong, conforming to the surrounding tissue. On average red calcophores are larger than the green ones, and green larger than blue, with mean diameters of 26.7 ± 2.3 µm, 23.8 ± 3.0 µm, 19.8 ± 2.8 µm. After isolation, calcophores take on a circular shape in projection due to internal pressure, revealing that they are membrane-bound.

#### 2. TEM

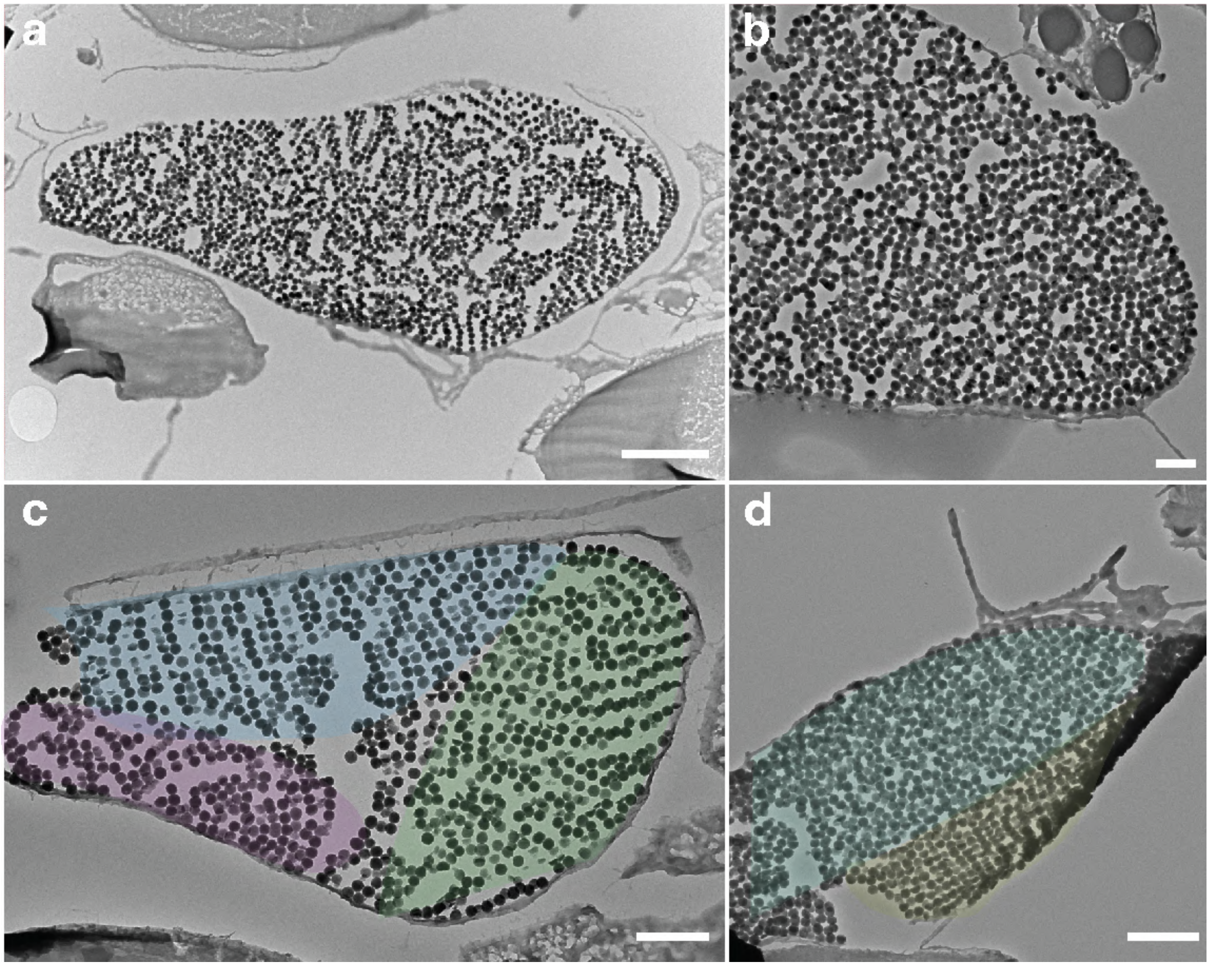


**Supplementary Figure 3**

a) Image of a cryo-cut TEM section of *Elysia viridis* calcophore, showing membrane bound nanoparticles with high electron density. Scale bar, 2 µm.

b) TEM image of *Elysia viridis* calcophore showing amorphous packing of nanoparticles, in a variety of 2D arrangements depending on the cut plane. Scale bar, 0.5 µm.

c) TEM image of *Elysia viridis* calcophore showing polycrystalline (colours represent discrete photonic crystal domains) packing of nanoparticles. Scale bar, 1 µm.

d) TEM image of *Elysia viridis* calcophore showing mixed amorphous and crystalline packing. Scale bar, 1 µm.

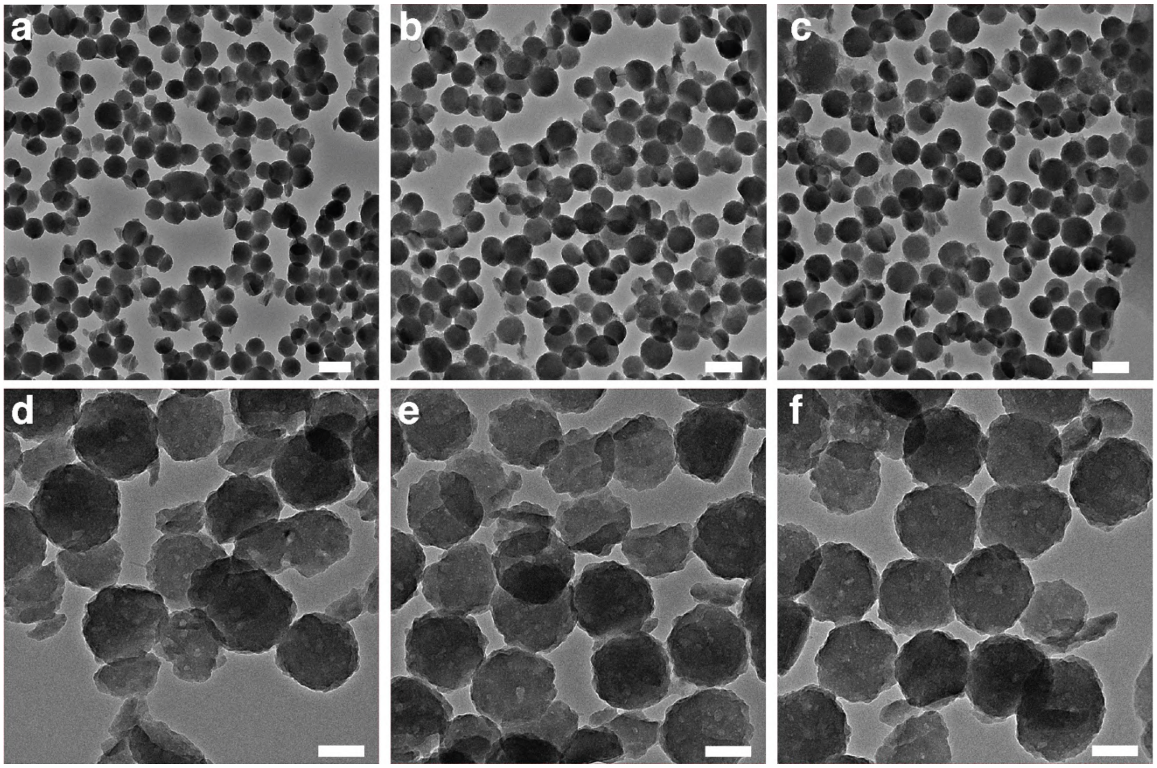


**Supplementary Figure 4**

TEM images of cryo-cut sections of *Elysia viridis* calcophore, showing spherical, rough nanoparticles. Some nanoparticles are broken during sectioning. Scale bars **a**-**c**, 200 nm, **d**-**f**, 100 nm.

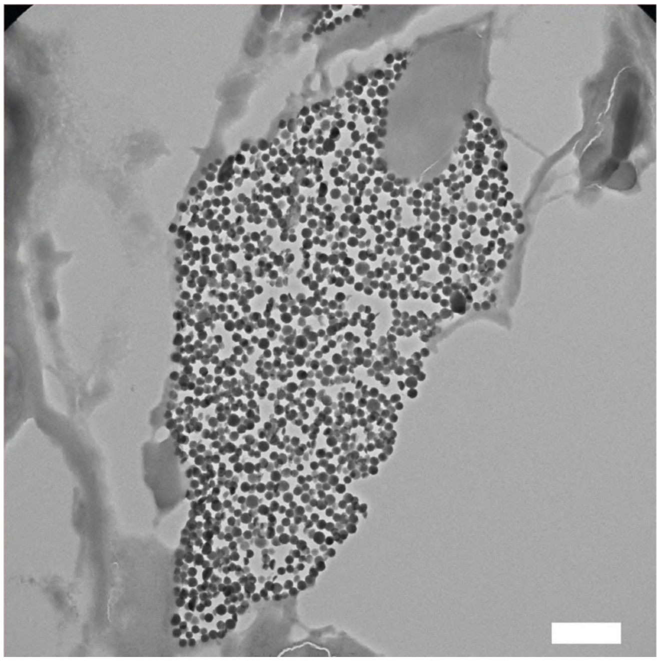

**Supplementary Figure 5**

Cryo-cut TEM section of *Elysia viridis* calcophore, showing the membrane bound nanoparticles with high electron density, and a cell nucleus located at the cell tip. Scale bar, 1 µm.

The mean nanoparticle diameter, measured over 20 calcophores, is 157 ± 25 nm, with maximum and minimum mean diameters of 216 ± 34 nm and 125 ± 16 nm respectively. The largest and smallest individual nanoparticles measured had diameters of 310 nm and 91 nm. The average polydispersity was 0.085, with maximum and minimum values for individual calcophores of 0.228 and 0.026.

#### 3. Optics

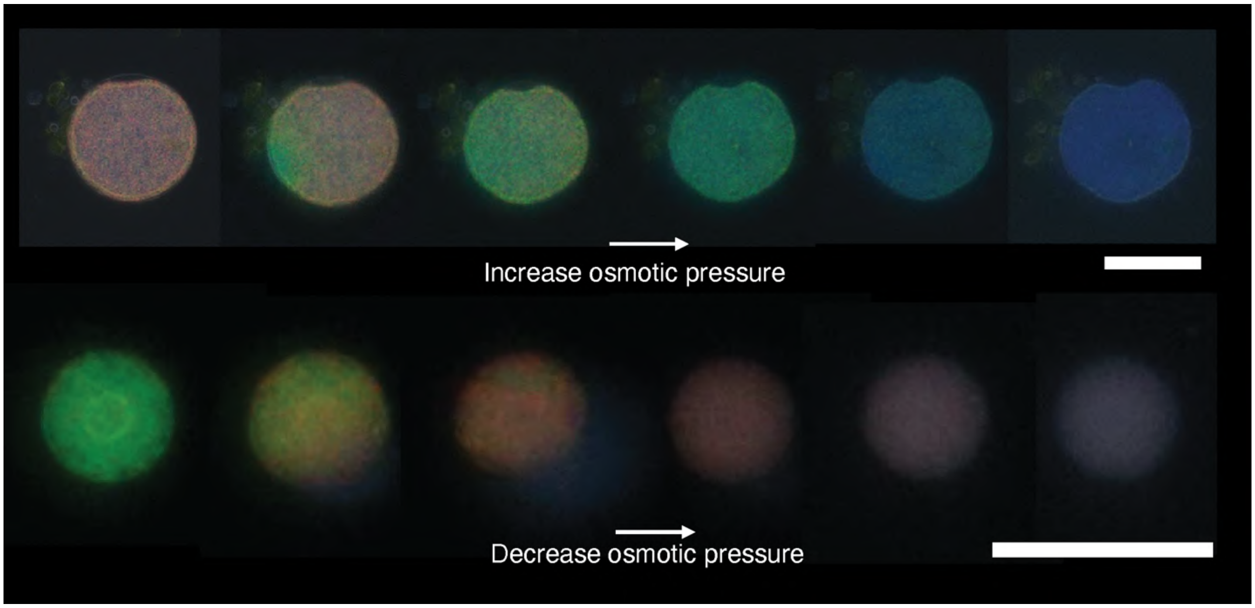


**Supplementary Figure 6**

Colour change of isolated calcophores under conditions of increasing (blue shift) and decreasing (red shift) osmotic pressure, induced by altering NaCl concentration. Scale bars, 25 µm (top). 20 µm (bottom).

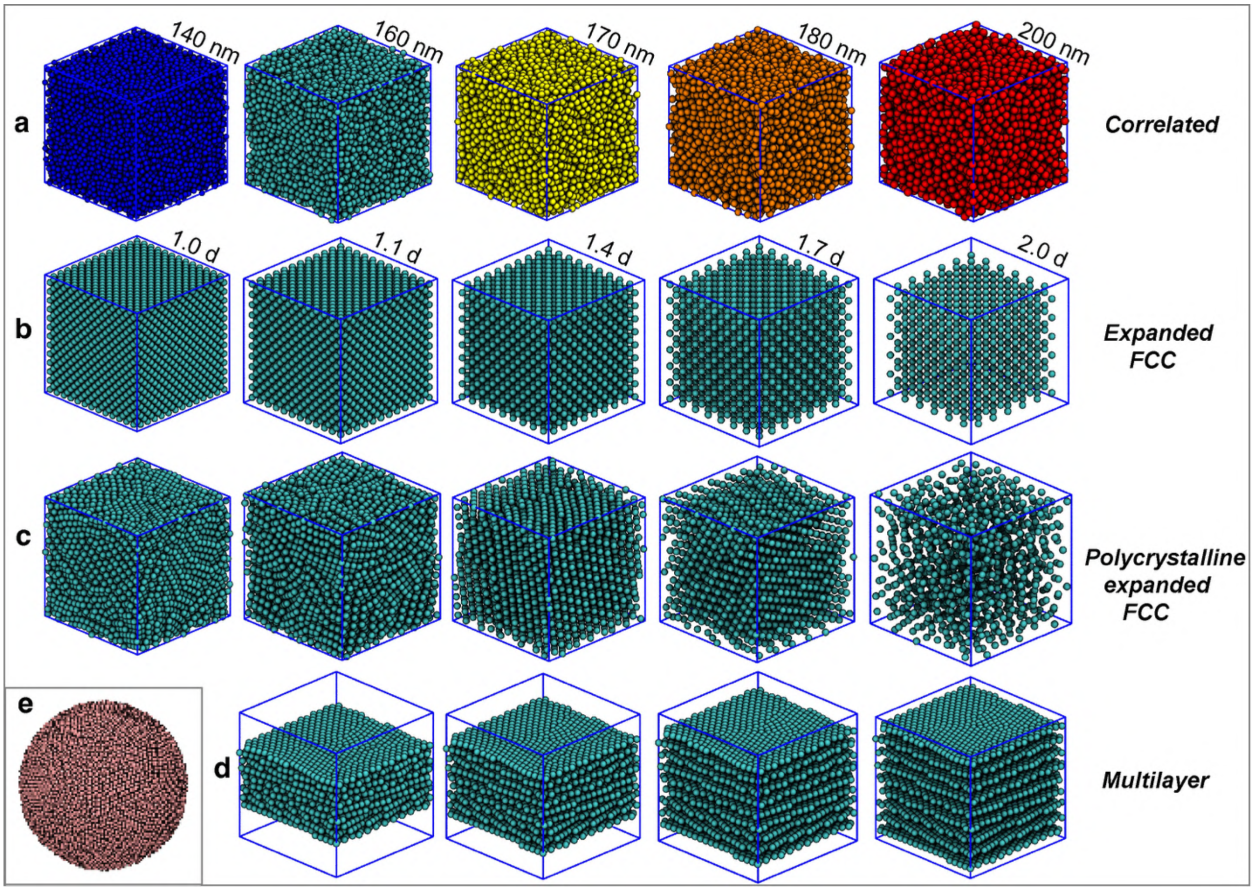


**Supplementary Figure 7**

Molecular dynamics simulations of the possible structures found in *Elysia viridis* photonic calcophores and used for FDTD simulations of optical reflectance.

a) Correlated structure, with particle sizes varying from 140 - 200 nm.
b) Expanded FCC photonic crystal with particle-to-particle distances varying from 1 d to 2 d (where d is the diameter of one particle, in this case 160 ± 5 nm). The (111) plane is perpendicular to the detector for simulations.
c) Same as b), but with polycrystallinity, i.e. the simulated structure is composed of multiple crystallographic volumes.
d) Multilayer stack structure where each layer is composed of spherical nanoparticles of diameter 160 ± 5 nm, and the plane spacing varies from 1.1 d to 2 d.
e) Representation of the spherical photonic unit used for FDTD simulations. Diameter = 5 µm.

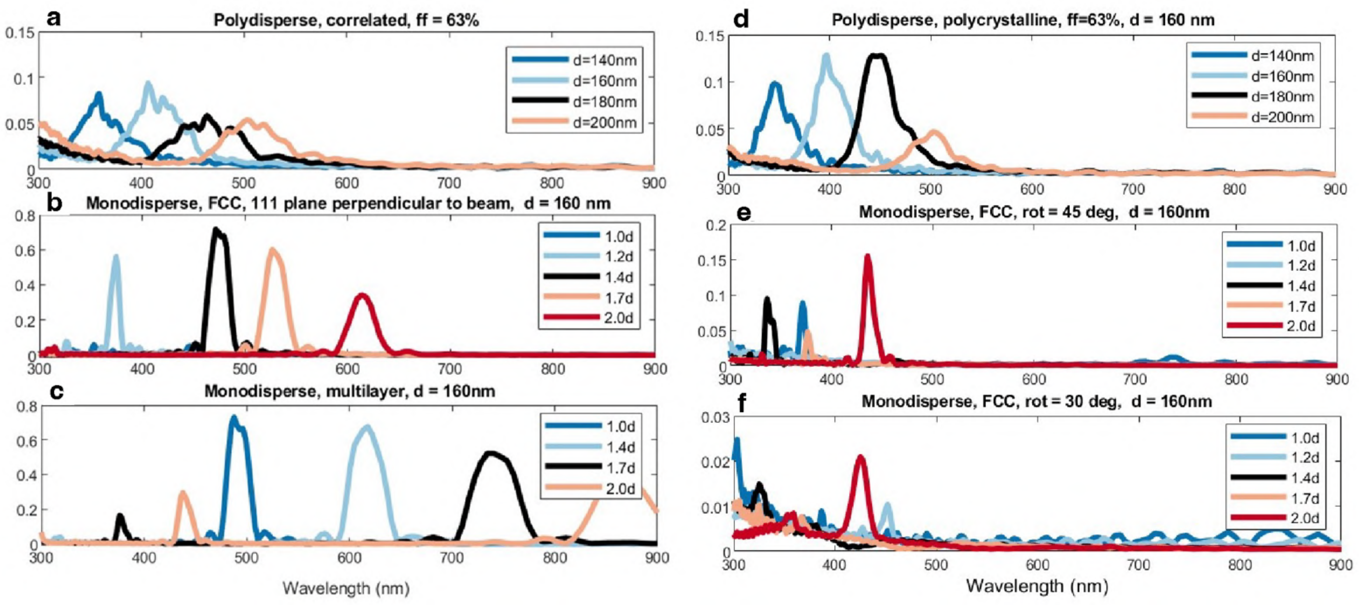


**Supplementary Figure 8**

FDTD simulations of 5 µm diameter spherical photonic structures found in *Elysia viridis*. A gaussian beam of radius 1.25 µm was used for all simulations. Simulations correspond to the structures in **Supplementary Figure 7**.

a) Simulated spectra for a correlated structure, with particle sizes varying from 140 - 200 nm.
b) Simulated spectra for an expanded FCC photonic crystal with particle-to-particle distances varying from 1 d to 2 d (where d is the diameter of one particle - in this case 160 ± 5 nm). The (111) plane is perpendicular to the detector for simulations.
c) Simulated spectra for a multilayer stack structure where each layer is composed of spherical nanoparticles of diameter 160 ± 5 nm and the plane spacing varies from 1.1 d to 2 d.
d) Simulated spectra for close-packed spheres with polycrystallinity, i.e. the simulated structure is composed of multiple crystallographic volumes for d = 140 - 200 nm.
e) Same as b) but with the structure oriented at 45 ° with respect to the incident beam.
f) Same as b) but with the structure oriented at 30 ° with respect to the incident beam.

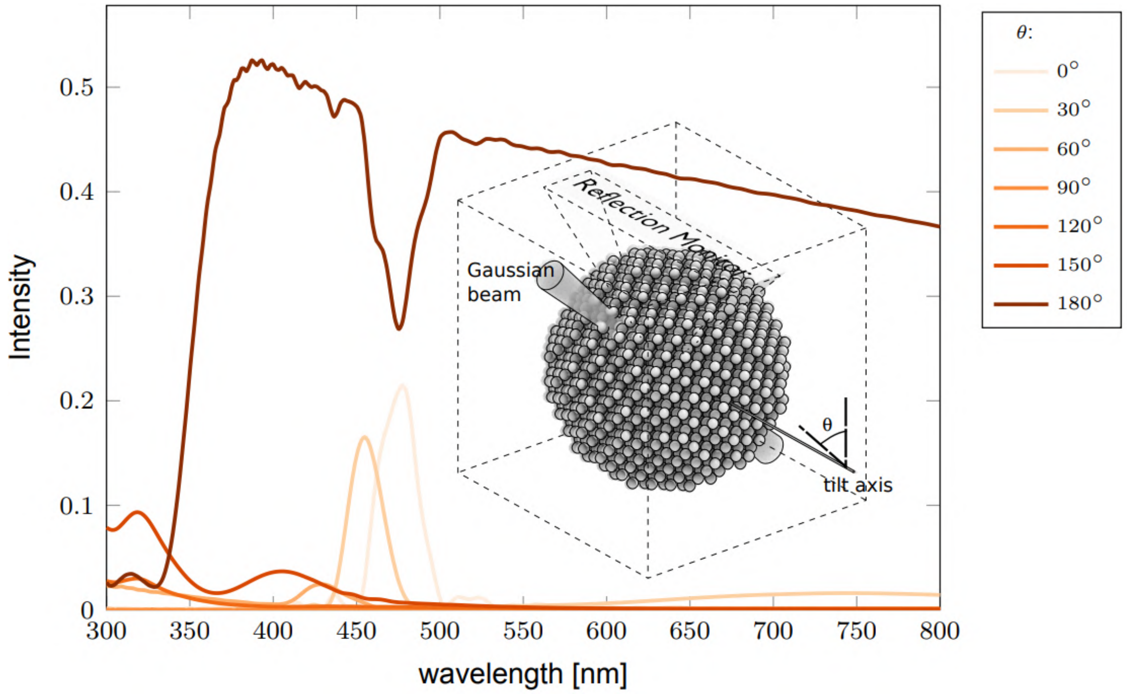


**Supplementary Figure 9**

FDTD simulation of the angular response of an expanded FCC photonic crystal. The incident beam is always perpendicular to the (111) plane, and the detector angle is varied. d = 170 nm, particle spacing = 1.4 d, filling fraction ~ 27.4%, Bounding sphere diameter = 3 µm. The reflectance peak height significantly drops at angles greater than 30 °.

Here we will elaborate in further detail on the simulations shown in **Figure 3** and show further simulations of alternate possible colloidal packing arrangements, including a multilayer (**Supplementary Figure 7d** and **Supplementary Figure 8c**) and FCC with illumination at an angle to the (111) plane. The first arrangement is a correlated system (**Figure 3d, Supplementary Figures 7a, 8a**). Here, we simulated particle diameters in the range 140 - 200 nm (as found through TEM), with a standard deviation of ± 5 nm for each particle diameter. As expected, due to the lack of long-range order, there is a photonic glass-like response, with broad, low-reflectance peaks and increasing peak wavelengths with increasing particle size.

We then simulated the expanded FCC structure with the (111) plane perpendicular to the beam direction (**Figure 3e, Supplementary Figure 7b, Supplementary Figure 8b**). This is the arrangement found through 3D cryo-FIB SEM imaging. For these (and the following) simulations, monodisperse spheres with a diameter of 160 nm were used. As shown in **Figure 3e (Supplementary Figure 7b)**, the simulated spectra are sharp, with peak reflectance of ~ 0.7 in the green region, and lower reflectance in the blue and red regions. These spectra closely match the experimental spectra for photonic crystals in **Figure 3h**. The red shift caused by increasing particle centre-to-centre distance corroborates the colour change seen upon evaporation in **Figure 3c** and **Supplementary Figure 6**, a 250 nm red shift is seen with a doubling of the centre-to-centre distance. In these simulations, the (111) plane is perpendicular to the beam, as expected in vivo because the (111) plane is the most densely packed, and because this arrangement minimises interfacial strain energy at the calcophore membrane. The polycrystalline (**Figure 3f, Supplementary Figures 7c, 8d**) structure (d = 140 - 200 nm) showed a response that resembles the halfway point between the expanded FCC and correlated structures. The reflectance peaks reach up to 15%.

**Figures 3g, h** show experimental spectra of individual calcophores. The narrow reflection peaks (up to 60% reflectance referenced to a silver mirror and a full-width half maximum of just 43 nm for the brightest spectrum) correspond to the expanded FCC simulated spectra in **Figure 3e** (**Supplementary Figure 8b**). The wavelength range of these reflectance peaks is from 470 - 695 nm, with the brightest calcophores in the green region. The bright (>20% reflectance) narrow peaks correspond to calcophores showing the crystalline appearance visible in **Figure 3**. The calcophores, which exhibit a more homogeneous appearance (photonic glasses), show distinct spectral properties (**Figure 3g, Supplementary Figure 8d**). The reflectance of these calcophores is typically below 20% (generally ~10%) and shows broadband reflectance. The red-most spectrum in **Figure 3g** (**Supplementary Figure 8d**) has a peak reflectance of 9.5% and a full-width half maximum of 120 nm. This broadband reflectance is typical of photonic glasses, where the light path is less well defined. In summary, this data confirms that the brightest domains within calcophores exhibit an expanded FCC colloidal packing, with the (111) plane perpendicular to the animal's surface; these domains are large enough to yield single-crystal-like responses. Another colloidal structure that could give very bright reflectance is a multilayer, where each layer is composed of close-packed particles. The optical response of this structure (beam perpendicular to the planes of the multilayer) was simulated, and bright reflectance of up to 70% was found (**Supplementary Figure 8c**). For some centre-to-centre distances, the simulated spectra of a multilayer system match the experimental spectra; however, no peaks were observed experimentally in the IR range, and EM data do not fit this structure, so we do not suggest it is present in vivo. Such a multilayer structure would require either specific, anisotropic interparticle forces or a highly controlled assembly mechanism, both of which are unlikely in this system. We also simulated the single crystal expanded FCC structure with the (111) plane tilted at 45 ° and 30 ° to the beam. The two simulations for expanded FCC structures at an angle to the beam show smaller reflectance peaks than the 0 ° simulation. This shows the angular dependence of reflection here and explains why there are dark spots/stripes in some regions of the photonic crystal-like calcophores in vivo. A more detailed simulation of this effect is visible in **Supplementary Figure 9**.

### 4. Interpreting calcophore distribution

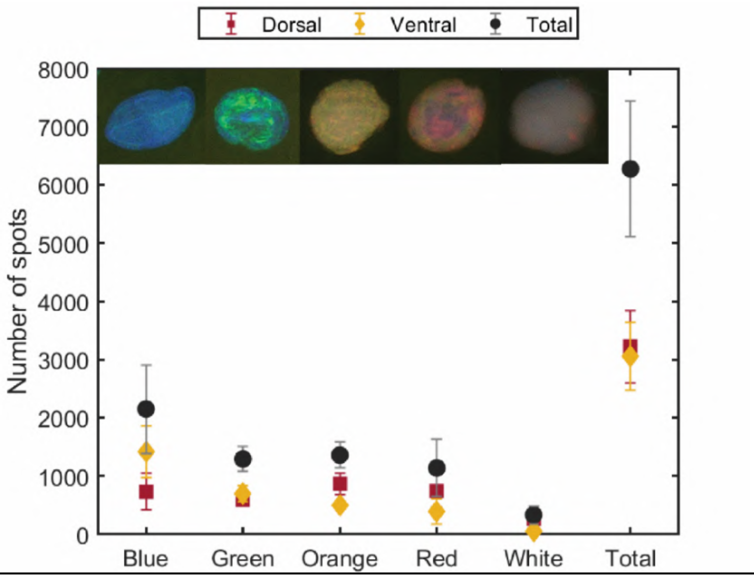


**Supplementary Figure 10**

Number of calcophores on the dorsal and ventral sides, and total for the colours blue, green, orange, red and white. Values are averaged over 6 individuals, and error bars represent standard deviation. Example Keyence microscope images of each colour are shown.

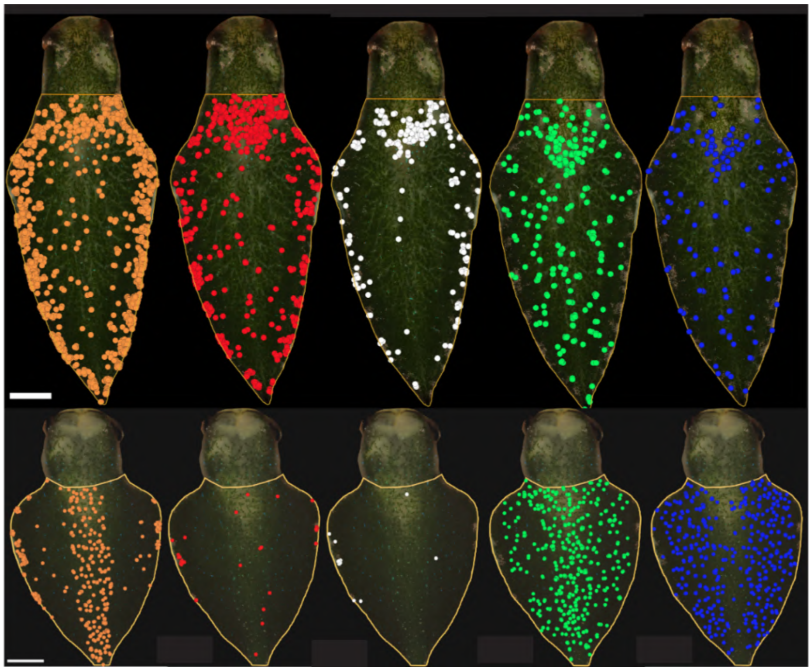


**Supplementary Figure 11**

Example images of the dorsal and ventral sides of an *Elysia viridis* individual. Orange, red, white, green and blue calcophores are mapped as coloured dots using the QGIS mapping software. Variation in pattern between colours is visible, such as the frequency of orange calcophores at the parapodial edge, or blue calcophores on the posterior side of the pericardial prominence. The head of the animal was mapped separately. Scale bars 0.5 mm.

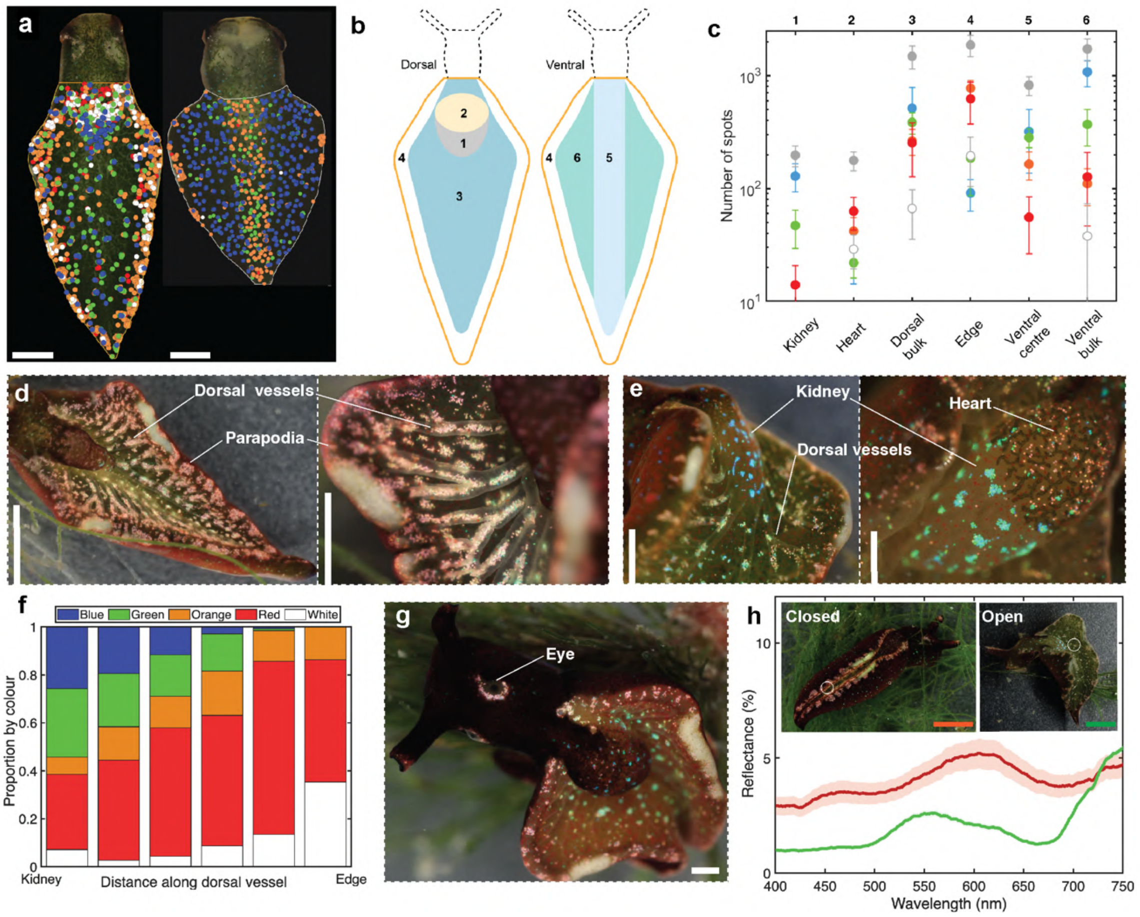


**Figure 4 / Supplementary Figure 12**

a) Example images of the dorsal and ventral sides of an *Elysia viridis* individual. Orange, red, white, green and blue calcophores are mapped as coloured dots using the QGIS mapping software. Variation in pattern between colours is visible, such as the frequency of orange calcophores at the parapodial edge, or blue calcophores on the posterior side of the pericardial prominence. The head of the animal was mapped separately. Scale bars, 0.5 mm.

b) 6 distinct regions defined by body mapping, these are: kidney (posterior pericardial prominence, area 1), heart (anterior pericardial prominence, area 2), dorsal bulk (area 3), edge (area 4), ventral centre (area 5), ventral bulk (area 6)

c) Colour mapping for the 6 regions defined in b) for the colours blue, green, orange, red, white and the total (marked in grey). Values are averaged over 6 individuals, and error bars represent standard deviation. Y axis has a log scale.

d) Digital microscope images of the dorsal surface of an individual, showing the clear correlation of dorsal vessels to calcophores, and high density of red and white calcophores at the edge of the parapodia. Scale bars, 2 mm, 0.5 mm.

e) Digital microscope images of the dorsal pericardial prominence (heart & kidney) of an individual, showing the high density of blue calcophore clusters above the kidney region, discrete red calcophores above the heart and red shift of calcophores along dorsal vessels away from the kidney towards the parapodial edge. Scale bars, 0.6 mm, 0.5 mm.

f) Histogram representing the proportion of each colour of calcophore along a dorsal vessel from the kidney to the parapodial edge (excluding the kidney and edge themselves). There is a clear red shift in calcophore colour along the vessels.

g) Example image of an individual with a clear ‘ring’ of red calcophores around the eye. Scale bar, 1 mm.

h) Double-ended probe measurements of an *Elysia viridis* individual with parapodia open and closed, showing the change in spectral response due to the photonic calcophores on the edge of the parapodia, which we suggest have a photoprotective role. Spectra are normalised to a silver mirror. The inset shows Keyence microscope images of *Elysia viridis* individuals in the closed (with a clearly visible strip of photonic calcophores) and open morphologies. Scale bars, 2 mm

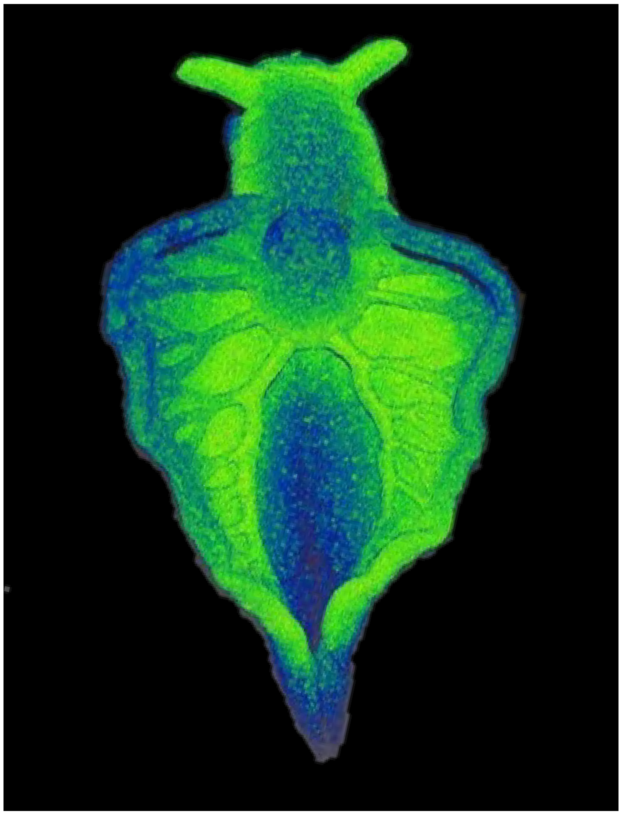

**Supplementary Figure 13**

Optical computed tomography (OCT) showing surface topography of a live, sedated adult *Elysia viridis* individual. The direct connection of the ramifying dorsal vessels to the kidney is visible.

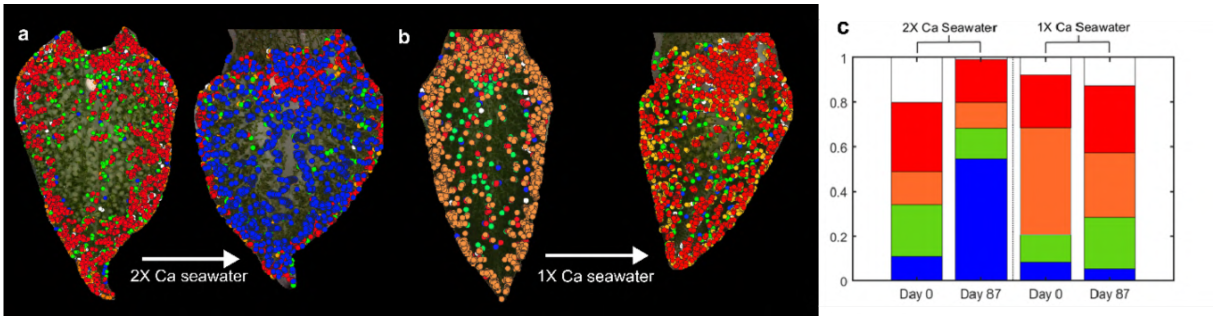


**Supplementary Figure 14**

a) QGIS point map of the dorsal side of an *Elysia viridis* individual 87 days after body regeneration in 2X Ca content seawater.

b) QGIS point map of the dorsal side of an *Elysia viridis* individual 87 days after body regeneration in 1X Ca content seawater.

c) Bar chart showing the numbers of each calcophore colour before and after regeneration in the two conditions above.

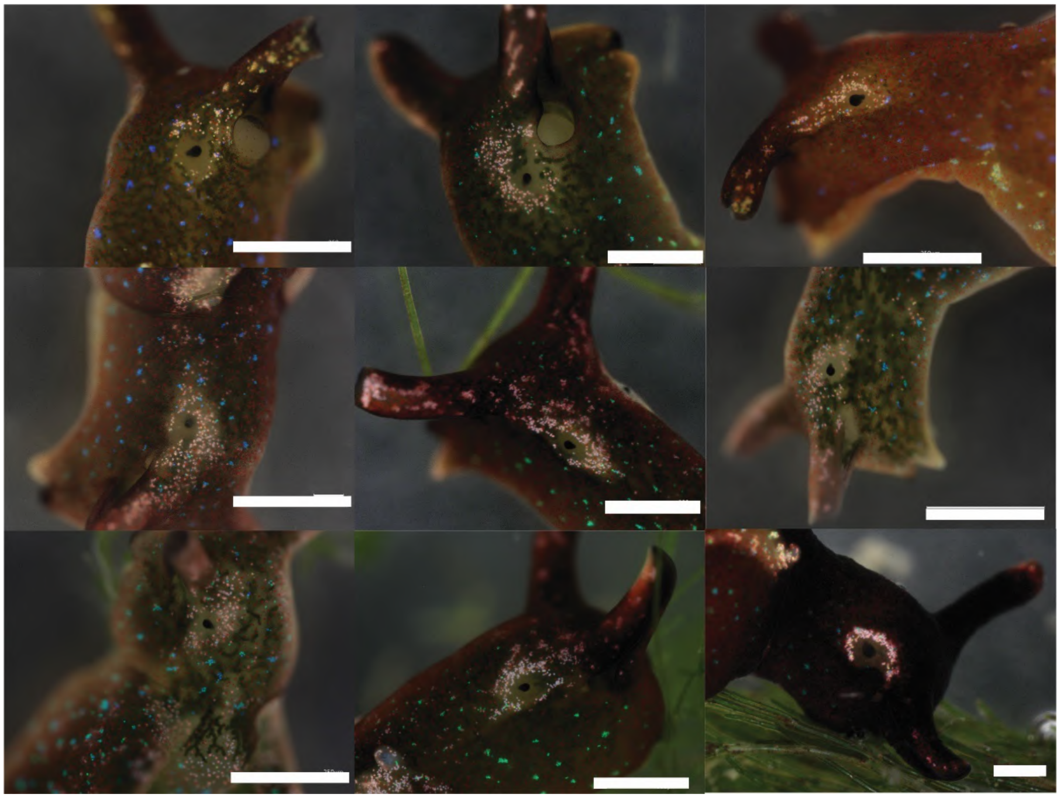

**Supplementary Figure 15**

Keyence microscope images of 9 *Elysia viridis* individuals showing rings of red calcophores around the eye. Scale bar, 500 µm.

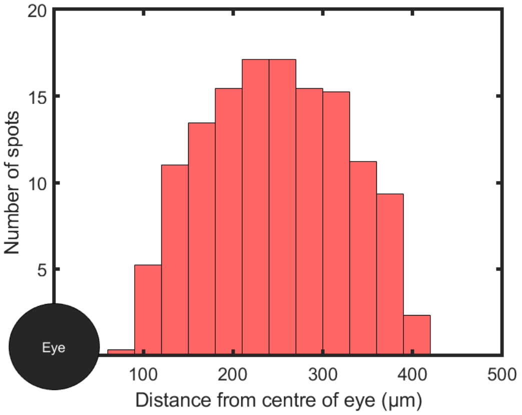


**Supplementary Figure 16**

Histogram of number of calcophores vs position from the centre of the eye (µm). A black circle is a scale representation of the eye. Distances were measured for 9 individuals (1 eye each).

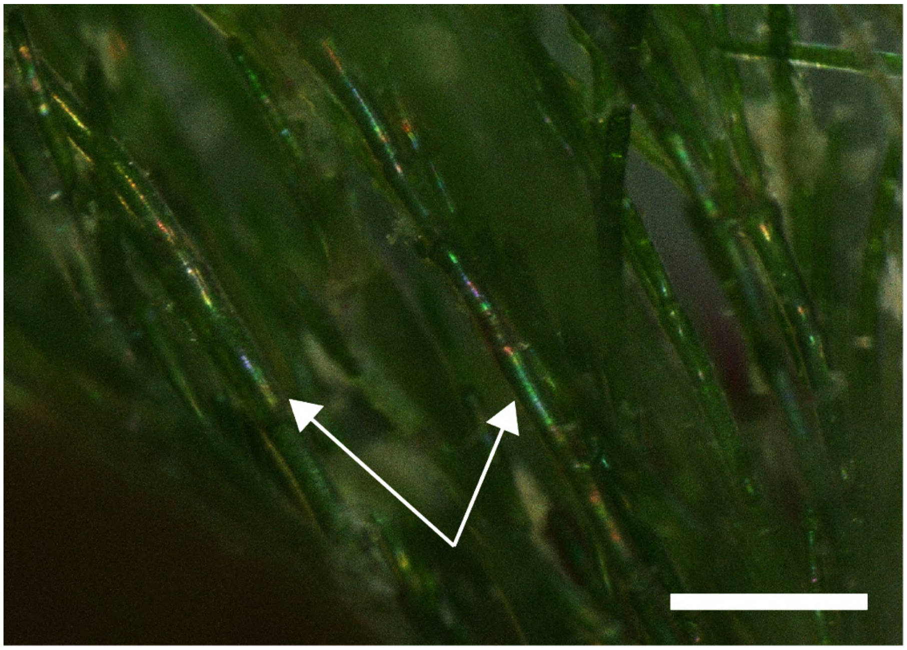

**Supplementary Figure 17**

*Cladophora rupestris* showing structural colouration (arrows). Scale bar, 500 µm.

The average total number of calcophores on the body of an individual slug is 6273 ± 1165, with 3220 ± 615 on the dorsal surface and 3054 ± 580 on the ventral surface. When separated by colour across the whole-body surface (**Supplementary Figures 10, 11**), we found that the highest proportion of calcophores is blue, a total of 2151 ± 759, followed by orange (1360 ± 219), green (1292 ± 216), red (1138 ± 490) and then white (333 ± 155). Colours show significant differences in their distribution, blue and green calcophores are mainly distributed on the ventral side (66% and 54% respectively) whilst orange red and white calcophores have a higher population on the dorsal side (64%, 66% and 82% respectively). Considering only the dorsal and ventral sides does not provide an accurate description of colour patterning, so the animal surface was partitioned into 6 distinct areas (**Figure 4b, c**). These areas were chosen based on the observations seen both in point maps, and from observation of individuals under the microscope. There are clear localisations of colours in particular areas of the animal such as the high concentration of clustered blue calcophores on the posterior side of the pericardial prominence (kidney, area 1), the dominance of individual red/white/orange calcophores on the anterior side of the pericardial prominence (heart, area 2), the high density of red/white/orange calcophores on the edge of the slug typically in clusters which bridge the dorsal and ventral sides (area 4). These patterns are shown in the example images in **Figure 4d, e**. The other patterns are less clear without point maps; a high density of clustered blue/green calcophores in the bulk of the surface area on both the dorsal and ventral sides (areas 3 and 6), and a strip of calcophores down the length of the ventral side, which is red-shifted from the calcophores in bulk (area 5).

Of the average number of 197 ± 41 calcophores in area 1 (the posterior side of the pericardial prominence, lying above the kidney), 65% were blue, and 89% were blue or green. The nearest neighbour index in this region was 0.57. This demonstrates that this area is dominated by blue and green calcophores, which form clusters (**Figure 4e**). In area 2 (the anterior side of the pericardial prominence, lying above the heart), the nearest neighbour index is 0.76, approximately 20% higher than in area 1 (i.e. less clustering). Of the average number of 177 ± 34 calcophores in this area 76% are red, white or orange demonstrating the expected pattern of discretely separated red/white/orange calcophores dominating this area. Area 3 (the bulk surface of the dorsal side) covers most of the dorsal surface of the animal and of the average of 1483 ± 348 calcophores in this area, 61% are blue/green. Area 4, corresponding to the edge of the parapodia, has the highest number of calcophores of the entire body, and the highest density by area. An average of 1869 ± 408 calcophores occupy this area, 85% of these are red, orange or white. As visible in **Figure 4d** the clusters here lie at the termini of the ramified dorsal vessels. The number of calcophores per length of the body in this region was an average of 196 ± 17 $mm^{-1}$, which (using the calculated average area of a red calcophore of 0.000543 $mm^2$) gives an average of 0.11 $mm^2$ covered by calcophores per mm length of the slug. In other words, this is the equivalent to each slug having a reflective strip which is 55 μm wide down the edge of each of its parapodia, an average total surface area of 1 $mm^2$ of edge calcophores per animal. The bulk of the surface on the ventral side (area 6) is again dominated by blue and green calcophores - of the 1722 ± 385 calcophores in this region 84% are blue or green. Region 5, the central strip of the ventral side of the slug (the 'foot') of the animal has an average total number of calcophores of 824 ± 155. There is a majority of blue and green calcophores here (73%), however, in this region, the dominant colour is always red-shifted from the dominant colour in region 6. If the dominant colour in region 6 is blue, it will be green in region 5, and if it is green in region 6, it will be orange in region 5.

In all individuals there is a red shift of calcophores from the kidney to the parapodial edges, and in individuals with an especially large number of calcophores, there is a clear correlation between the position of the calcophores on the dorsal side and the dorsal vessels, which protrude from this surface (**Figure 4d, e**). These vessels originate at the pericardial prominence, ramifying across the surface. This has also been reported in other species of sacoglossa[1]. In individuals where this effect is clear, following the vessel away from the kidney to the edge of the animal the colour of calcophores red-shifts (**Figure 4e**). For individuals where the density of calcophores is not high enough to trace a continuous line under the vessels, the colour shift is still clear (**Figure 4g**). Because blue calcophores have a higher density of nanoparticles than red ones, the mineralization process may be linked to their colour distribution. In other words, the high density of calcophores at the kidney implies a high degree of mineralization in this region, resulting in calcophores with a high filling fraction and therefore photonic crystals which reflect blue light. The connection of dorsal vessels to the kidney region (**Supplementary Figure 13**) and to calcophore positions suggests that they may act as transport vessels for mineralization precursors, whilst the red shift in calcophores along their length towards the clusters at the termini of these vessels demonstrates a tend to calcophores with a lower filling fraction. This is the result of a lower rate of mineralisation with decreasing proximity to the kidney. We therefore suggest that these features of the calcophore colour distribution are linked to formation. To confirm that colour is linked to the rate of biomineralization (which we believe to be related to the internal concentration of calcium ions, since calcophores are composed of mineralised calcium nanoparticles) we used body regeneration[2] as a proxy for initial body growth and calcophore formation, see **Supplementary Figure 14**. A slug was mapped before and after regeneration in seawater with doubled calcium concentration (further increase calcium led to the death of the animals). As visible in **Supplementary Figure 14**, the body of the slug grew back with a strikingly larger number of blue calcophores with large clusters across the dorsal surface, far more than had been seen in any wild-caught samples. This was then compared to a slug which had regenerated its body in normal seawater, to determine whether it was the calcium concentration making the difference. To quantify the effect, the calcophores were counted before decapitation and 87 days afterwards. We observed that the population of blue calcophores increased from 11% to 55%, a 5-fold increase. When compared to the control slug that regenerated its body in normal seawater, for which we observed a 3% decrease in proportion of blue calcophores from 8% to 5% the difference is striking. The slug grown in 2X calcium seawater slug had a drop in the proportion of red/orange/white calcophores on its dorsal surface of over 50%, whilst the control slug remained within 8% of its original proportion. It is clear from **Supplementary Figure 14** that the greatest colour change comes in the increase in the number of blue calcophores. So, the availability of calcium in the water of a growing slug may have a large effect on the colour of an individual's calcophores, by increasing the rate of mineralization and therefore the final nanoparticle packing density.

Whilst we suggest the colour and position of calcophores are intimately linked to their biogenesis, we also believe that their distribution is functional. Heads were mapped separately from the body. Images of the slugs were taken whilst the individuals were free roaming and naturally positioned themselves eye-to-eye with the microscope (**Supplementary Figure 15**). In every individual, there is a 'halo' of calcophores around the eye, which are entirely red/white. The average distance from the centre of the eye to a calcophore (the radius of the halo) was 248 ± 15 µm, and the average radius of the eye was 51 ± 8 µm, such that on average a calcophore in this ring lies 196 ± 11 µm from the edge of the eye (**Supplementary Figure 16**). The average eye has 134 ± 13 calcophores. Given that this is consistent across every individual studied, we suggest that it may be functional, acting to either augment vision by increasing the amount of light reaching the eye via scattering from the ring of calcophores or increasing the conspicuousness of the animal's head for orientation during mating. Sea slugs do not have complex eyes and are thought to only detect changes in local light intensity[3]. For this reason, the specific colour of calcophores in different regions cannot be attributed to this function. However, the locations of bright regions of calcophores, such as above the kidney or around the eye, could increase their conspicuousness to other slugs, perhaps helping them orient during mating. The red calcophores along the parapodia could help outline the body and aid in locating other individuals.

The high density of calcophores at the parapodial edges of *Elysia viridis* is consistent across every individual. Kleptoplasts inside the cells of *Elysia viridis* are highly vulnerable to light damage, and it has been reported in multiple species, including *Elysia viridis,* that animals will close their parapodia in response to high light[4,5]. Not only does the closing of parapodia reduce the overall surface area of the animal, thereby reducing light exposure to chloroplasts, but it also directs the clusters of calcophores on the parapodial edges upwards. If this region of the animal is approximately 1 cm long, then the result is roughly 1 $mm^2$ of area covered by reflective calcophores. With an average of 20% reflectance for the red calcophores and a maximum irradiance of 1 $kWm^{-2}$ or 0.001 $Wmm^{-2}$ in Bretagne during summer, the calcophores could actively shield a slug from 0.2 mW of solar irradiation. This could be a key factor, amongst others, which allows this species to extend chloroplast lifetime. To summarise, the closing of parapodia under high light will direct a reflective photonic shield of calcophores towards the light, meaning that a proportion of the light striking the remaining surface area will be backscattered, as an active photoprotection mechanism. The calcophores are therefore concentrated here so that the animal can selectively decide when to limit light to the chloroplasts. In low-light conditions, maximising solar irradiance is key for chloroplasts. To quantify this effect, we obtained reflectance spectra at open and closed parapodial positions. **Figure 4h** shows spectra taken of an individual with parapodia open and closed, showing the reflective peak in the red region in the closed position, covering the chlorophyll spectral signature. **Figure 4h inset** shows example images of individuals in the open and closed morphologies.

Structural colour often has a communication function, such as aposematism in nudibranchs[6], or sexual signaling. In the case of *Elysia viridis* (even though they contain toxic secondary metabolites[7]), there is unlikely to be an aposematic function due to the sparse nature of the colour here. The calcophores might resemble the reflection from the structural colouration of the algae that they live in symbiosis with, increasing the camouflage of the slug in its environment, see **Supplementary Figure 17**. However, the degree of conspicuousness imparted by structural colour is very slight and inconsistent between individuals, whilst for other sacaglossa such as those in the *Thuridilla* genus this is a likely function[8].

Healthy *Elysia viridis* individuals are green and therefore camouflage well with the green algae which they eat, as is true for many sacoglossa[9]. At least 3 of the species of green algae which they eat also show structural colour,  these are *Bryopsis hypnoides*, *Cladophora rupestris* (**Supplementary figure 17**) and *Valonia ventricose*. It is possible then that the structural colour in *Elyisa viridis* is a form of masquerading. Camouflage in sea slugs is common, and some species of sacoglossa are known to use ventral colouration to masquerade as sand, despite the green appearance with open parapodia[10]. Although the verdant appearance of sacoglossa provides an overall background match, its structural colour could offer disruptive camouflage. In *Elysia viridis,* the outer surface of the animal is made more conspicuous under the microscope by the high density of photonic calcophores along the edges of the parapodia. However, patches may act to blur the edge or confuse the outline of the animal[11] or act as a form of disruptive camouflage[12].  In other words, the iridescent calcophores in *Elysia viridis* could distract from the overall shape of the animal by increasing the visual noise[12–14]. It is also possible that calcophores are not strictly functioning as colourants but only act as calcium sinks for disposal. Many low-metabolic-rate molluscs are thought to accumulate calcium concretions over the course of their short lives, which do not constitute a significant burden[15].

#### 5. Structurally coloured reproductive organ

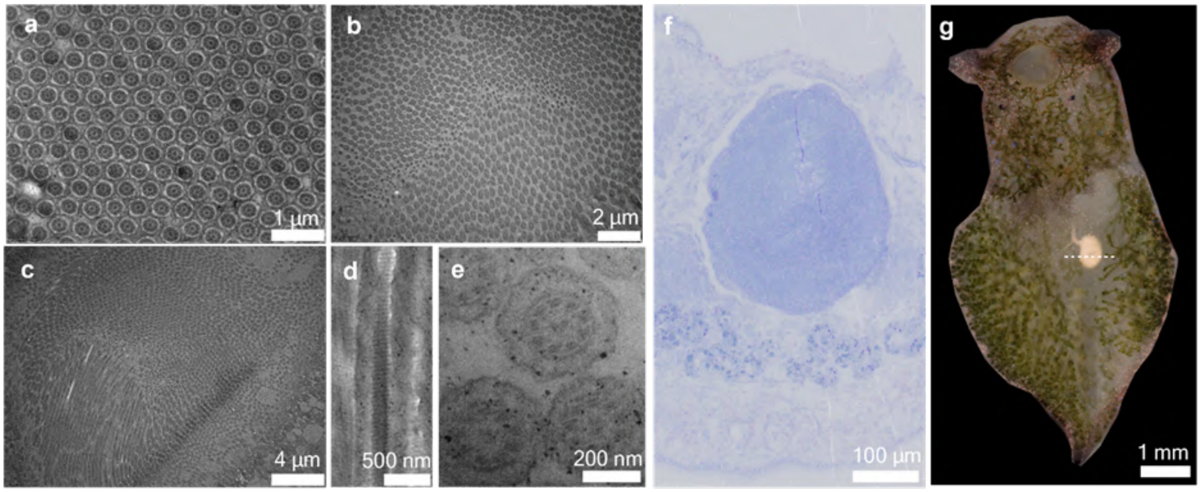


**Supplementary Figure 18**

a) TEM image of hexagonally packed 2D photonic crystal of *Elysia viridis* sperm.

b),c) TEM images showing a wider view of the reproductive organ, with individual motifs changing from circular, to longitudinal depending on their orientation relative to the cut surface.

d) Cross-sectional view (longitudinal) showing 'paracrystalline' layers.
e) Cross-sectional view (transverse) showing 9 coarse fibres surrounding an axoneme.
f) Bright field microscope image of a section stained with toluidine blue showing reproductive organ (blue), posterior to the pericardial prominence.
g) Keyence microscope image of an *Elysia viridis* individual showing the pink/orange reproductive organ and the cross section viewed here in TEM.

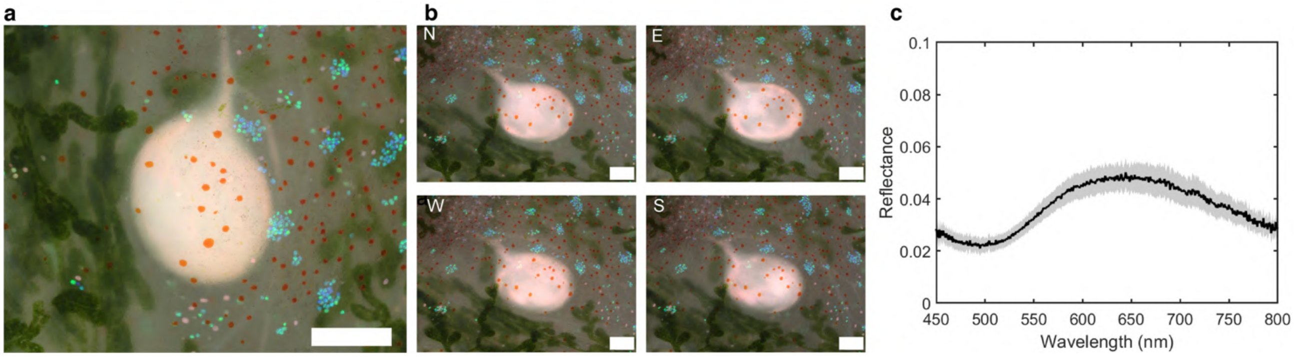


**Supplementary Figure 19**

a) Keyence microscope image showing the glowing orb. Scale bar, 200 µm.
b) Keyence microscope images varying the incident light direction (north, east, south, west). Scale bar, 200 µm.
c) Averaged reflectance spectra of 20 positions across a structurally coloured *Elysia viridis* reproductive organ showing broadband reflectance with a peak at ~640 nm. Shaded error bars represent standard deviation.

When taking TEM sections of a slug sample which had been fixed and osmicated, a large area lying in the centre of the animal, to the posterior side of the pericardial prominence, was found to be composed of a distinct ultrastructure. The size and position of this organ in TEM corresponds to that of the ‘glowing orb’ which had previously been imaged with light microscopy, but the photonic structure and functional significance of this ‘organ’ were yet unknown, and unreported in the literature. The TEM images of different sections in this region show what appears to be elongated tubes or villi-like cells. In many regions of the organ, these cells were highly aligned, with widths on the order of 200 - 300 nm, a suitable scale for orange/red structural colour. It should also be noted that this organ is not visible in juvenile animals, which may suggest its reproductive function. Reid et al.[16] describe ultrastructural details of cells in *Elysia maoria,* which resemble **Supplementary Figure 18**, assigning the cells as sperm (with size varying depending on whether it is a spermatocyte, or a mature sperm). Further literature searches reveal that the cells found here highly resemble the ultrastructure of sperm cells in other gastropods. Healy et al.[17] describe ‘Complex mitochondrial derivative sheathing the axoneme and usually nine coarse fibres; no unmodified cristae; inner and outer paracrystalline layers usually present’. These 9 coarse fibres surrounding the axoneme are visible in **Supplementary Figure 18e,** whilst the paracrystalline layers are visible in **Supplementary Figure 18d**. Due to refractive index contrast and their high alignment in certain areas, the result is a 2-dimensional photonic crystal (hexagonal packing is visible in **Supplementary Figure 18a**). The alignment varies across the volume and resembles a ‘wave-like’ structure, similar to long grass blowing in the wind. This is reflected in the optical appearance of the organ.

**Supplementary Figure 19a** shows a Keyence microscope image of the region described above. Compared with the calcophores, the colour is more diffuse; however, it appears somewhat opalescent and brighter than the orange carotenoid spots above and surrounding it. To confirm that this is indeed structural colour, modulating the incident light in 4 directions (north, east, south, west) reveals that this structure is iridescent. In other words, the direction of the incident light changes the optical appearance of the structure. Reflectance spectra show a broadband reflectance peak centred at 640 nm (**Supplementary Figure 19c**), with a full width at half maximum of ~ 200 nm and peak reflectance of 5%. As shown in the TEM images of this structure, the structure is a 2D hexagonal photonic crystal in which the repeating unit of the photonic crystal is a sperm cell. There is a very high degree of order in some regions, but reflectance remains small. We suggest that this is due to a low refractive index contrast between the sperm and its surrounding medium.

Unlike for calcium carbonate, these cells are composed of organic materials, likely lipids and proteins. What’s more, much of the structure shows little ordering. The photonic unit has a well-defined size (since the sperm cells are consistent in size), so it is the disorder and the lack of high-index contrast that give this organ its diffuse orange/pink appearance. This structurally coloured reproductive organ is large enough to potentially be detectable to other slugs. Perhaps a higher sperm count leads to a larger glowing orb, and healthy sperm align into more perfect photonic crystals, appearing brighter. Therefore, the function of structural colour in this organ could be to signal to other slugs an individual's reproductive fitness. This structure has not been reported for *Elysia viridis* and is the first reported example of a photonic crystal whose repeating unit is an animal cell.

#### 6. Calcophore formation

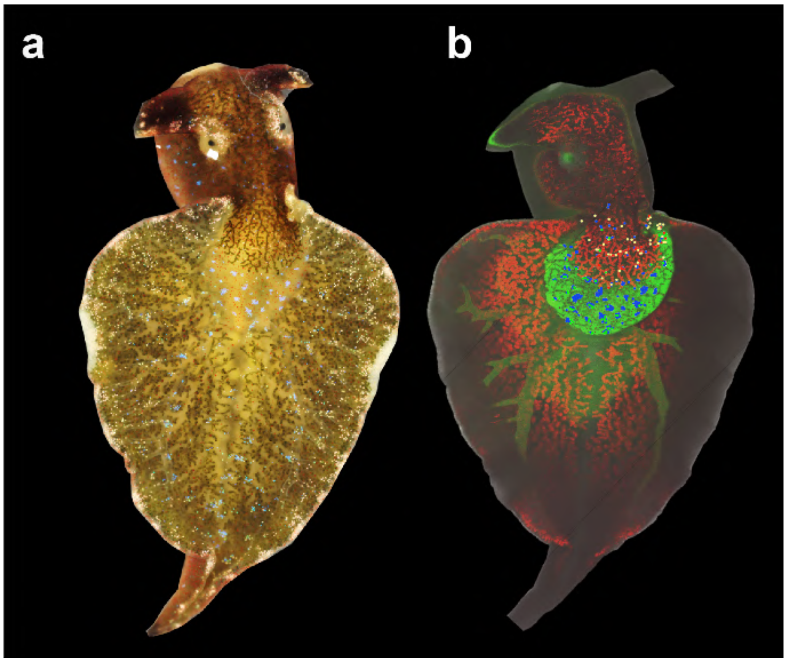


**Supplementary Figure 20**

a) Keyence microscope image of the same *Elysia viridis* individual used in whole body confocal fluorescence imaging, one hour later.

b) Superimposed mapping of calcophores above the pericardial prominence, showing the perfect correlation between the position of blue calcophores and the calcium-dense kidney. Calcophores were mapped using QGIS. Red is chlorophyll autofluorescence, green is calcein fluorescence.

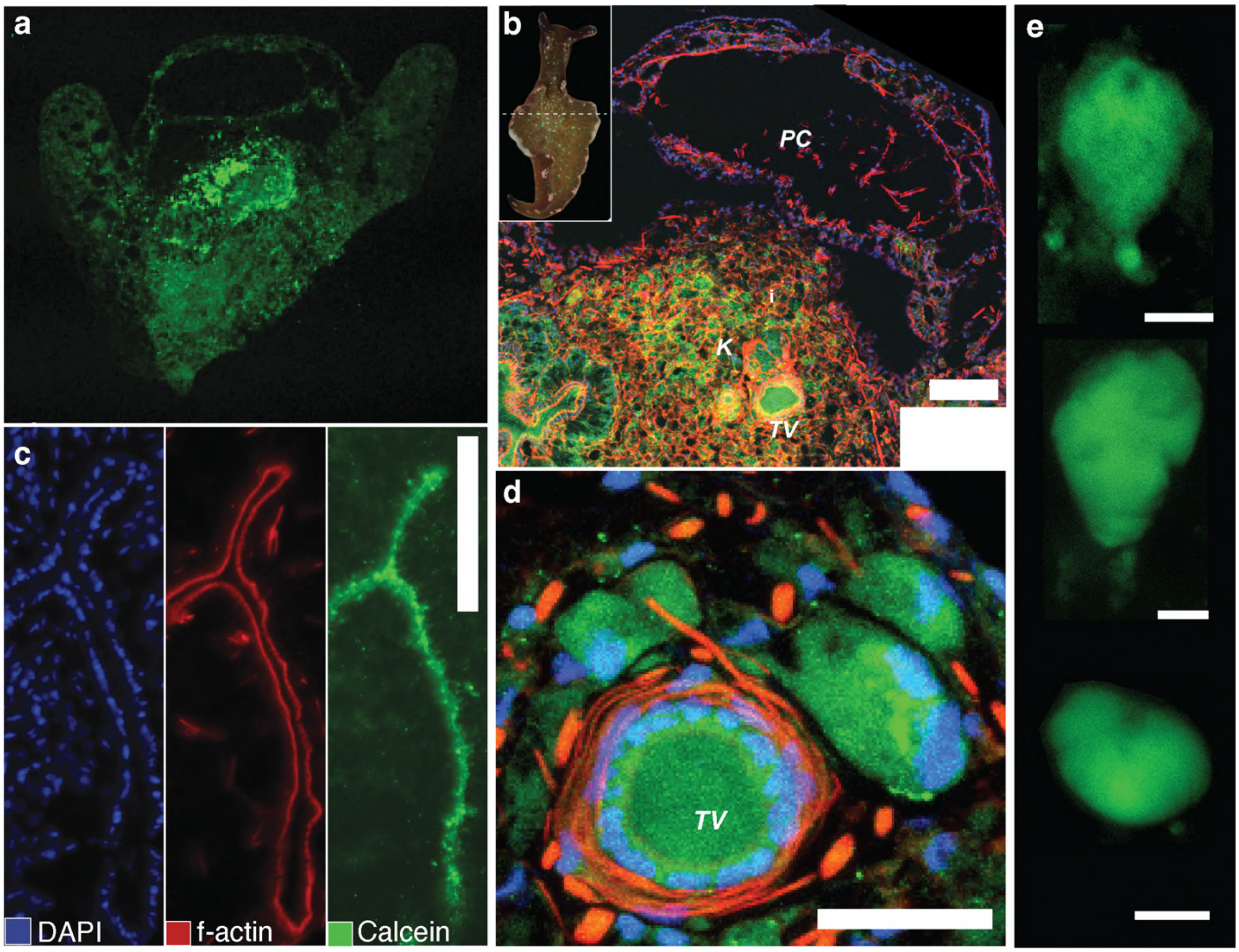


**Supplementary Figure 21**

a) 12 µm thick fixed *Elysia viridis* section of the kidney/heart and parapodia, showing dense calcein fluorescence (green) in the kidney region.

b) High magnification image of a 12 µm thick fixed *Elysia viridis* section of the kidney/heart, showing inhomogeneous distribution of calcein fluorescence (green). DAPI (blue) and f-actin (red) are also visible. Scale bar, 100 µm. Inset (top left) shows the position of the section in the body. The 2 heart cavities above the calcium-rich kidney are visible (the cross-section of a vessel transporting calcium is visible on the right side of the kidney).

c) Immunofluorescence of a 12 µm thick fixed *Elysia viridis* section. Stained with DAPI, phalloidin and calcein (blue, red & green fluorescence) demonstrating calcium-containing internal transport vessels. Scale bar, 100 µm.

d) Cross-section of a transport vessel in the parapodia, connected to calcified bodies [we believe to be calcophores] via a smaller vessel. Scale bar, 100 µm. K = kidney, PC = pericardium, TV = transport vessel. Scale bar, 25 µm

e) Live confocal imaging (maximum intensity projections) of possible calcophore precursors showing conduit-like extension with calcein fluorescence at the distal end to a dark region (which may be the cell nucleus). Internal fluorescence texture is smooth compared to that of a fully formed calcophore in **Figure 3c**, implying the calcium in these cells may be in the form of a dense liquid precursor. Scale bar, 10 µm.

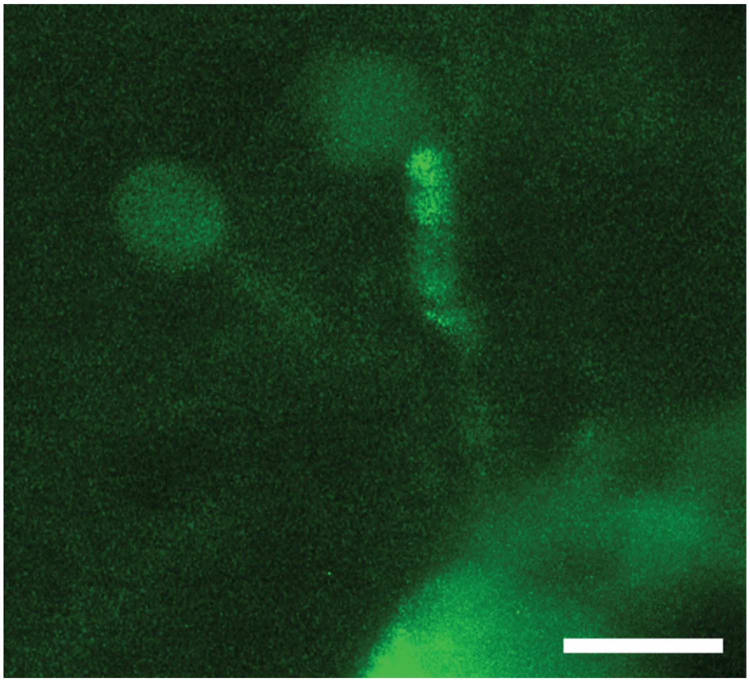

**Supplementary Figure 22**

Live confocal image (Zeiss LSM 980 with Airyscan 2 20X magnification) of an *Elysia viridis* individual showing two calcium dense regions, both connected to a larger vessel via small vessels. Region of interest is close to the parapodial edge. Scale bar, 10 µm.

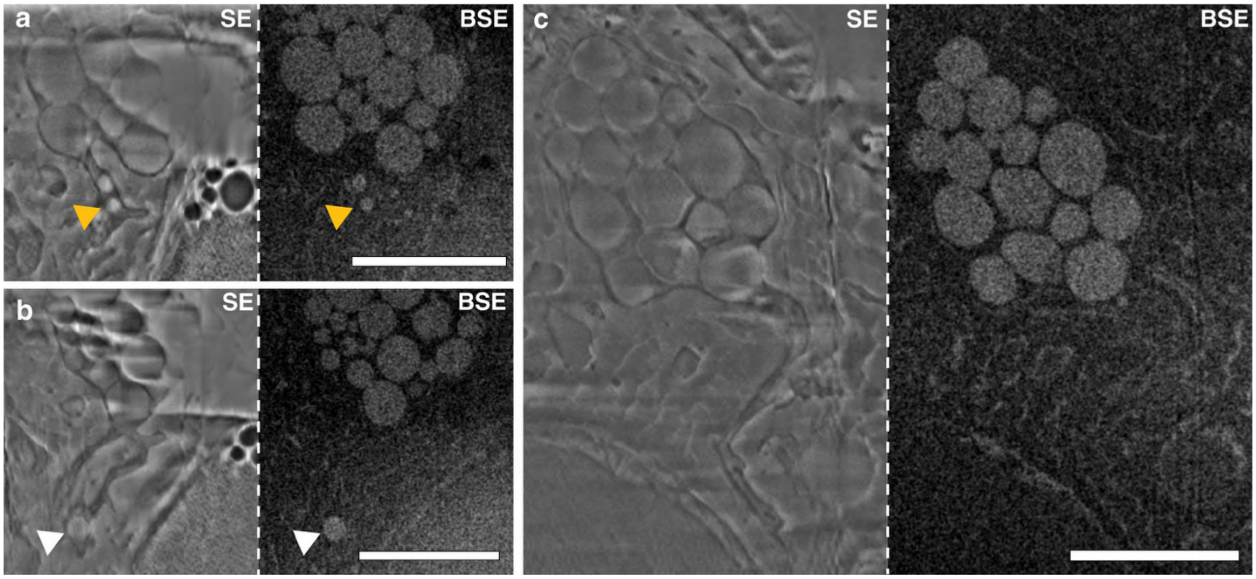


**Supplementary Figure 23**

2D Cryo-FIB secondary electron (SE) and backscattered electron (BSE) images, showing three regions of interest (**a**, **b** and **c** [**a** and **b** show images of different planes of the same developing calcophore]). Spherical bodies are visible within a membrane-bound volume that has a tube-like appendage or 'conduit' emerging from one end. The images show bodies located within this conduit, perhaps during transport. Yellow and white arrows highlight spherical bodies located inside the calcophore conduits and correspond to the yellow and white arrows in **Figure 5g, h.** Scale bars, 5 µm.

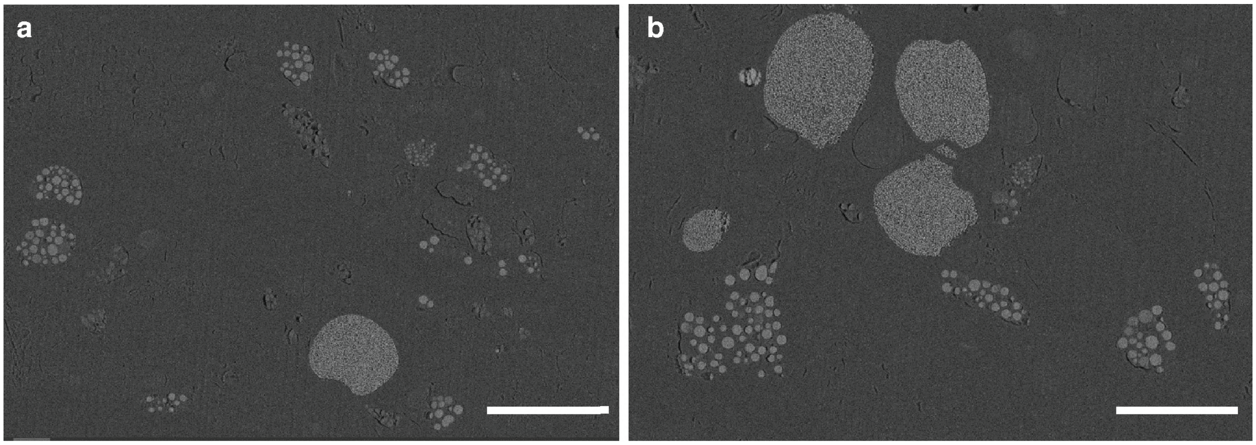


**Supplementary Figure 24**

a), b) Two examples of block face SEM in BSE imaging mode of freeze-substituted and embedded dorsal surface tissue, showing calcophores in close proximity to clusters of high BSE bodies. Scale bar, 20 µm.

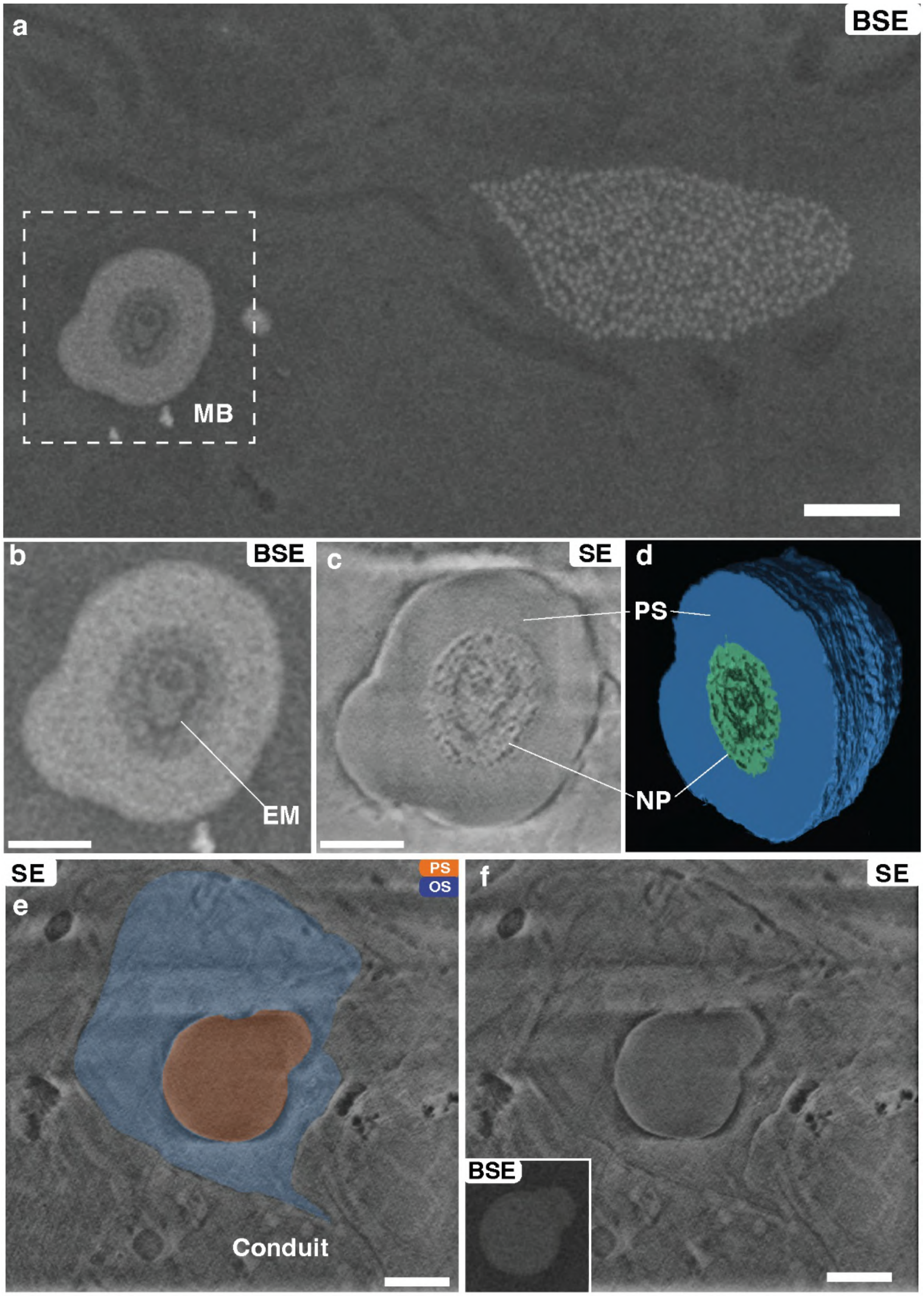
a
BSE
MB
b
BSE
EM
c
SE
d
PS
NP
SE
e
PS
OS
Conduit
f
SE
BSE

**Supplementary Figure 25**

a) Backscattered electron (BSE) SEM images taken using serial FIB-SEM recording of a mineralizing body found in close proximity to a calcophore. The mineralising body is found at the same sample depth, 10 µm from the fully formed calcophore. Scale bar, 2 µm. MB = mineralising body

b) BSE image taken using serial FIB-SEM imaging of a mineralising body. The body is located ~ 10 µm from a fully formed calcophore (at the same depth. Four concentric rings of low and high signal are visible, surrounded by a homogenous band of high signal. We suggest that the rings may represent cycles of exclusion and mineralisation of water and metal ions. EM = excluded material, PS = precursor solution, NP = nanoparticles

c) Secondary electron (SE) image of the same material in e). Concentric rings are visible here, with those corresponding to the high BSE signal showing nanoparticle-like texture; this texture corresponds to the texture in the high signal BSE bands. The outer band here is homogenous, implying an amorphous or liquid-like mineral state. Scale bar, 1 µm.

d) 3D FIB-SEM reconstruction of sequential backscattered images of the mineralising body in e). The low BSE signal volume has been removed, the high BSE, nanoparticle-like signal is coloured green, and the homogenous high BSE band is coloured blue, showing the formation of nanoparticles from the centre of the body.

e) SE SEM images taken using serial FIB-SEM recording of a mineralising body found in close proximity to a calcophore. The images shown here are close to the end of the backscattered volume shown in **Figure 5**. Using secondary electron imaging, the mineralising body is visible at the centre of the volume (coloured orange, and labelled PS [precursor solution]). The mineralising body is surrounded by an outer bulk volume (coloured blue and labelled OS (outer solution), with a small vessel extending from one end. We propose that this is a calcium transport vessel. The inset shows the backscattered signal corresponding to the precursor solution highlighted in orange, demonstrating its high mineral content. Scale bar, 1 µm.

f) Region of interest from e) with an inset showing the backscattered signal.

To examine the formation of calcophore during the process of mineralisation, we imaged an area containing calcophores using cryo-FIB SEM. We observed a spheroidal mineralising body (**Supplementary Figure 25)**, maximum diameter 5 µm [this truncated volume is an underestimate of the true diameter]) enclosed within a larger membrane (maximum diameter 10 µm) with a small vessel extending from one end. **Supplementary Figure 25** shows the relation of this mineralising body to a calcophore located 10 µm away from the edge of the calcophore, at the same depth. The bulk outer layer of the mineralising body is homogeneous in secondary electron (SE) mode, with a high signal in back-scattered electron (BSE) mode, suggesting a high metal-ion content. The homogenous, unfaceted outer band suggests these ions are in an amorphous state. Wolf et al.[18] showed that $CaCO_3$ mineralisation from a dense liquid precursor in levitated droplets can occur in the core rather than at the boundary of a droplet. In this study, opalescence emerges from nanoparticle aggregates expanding outwards from the droplet centre. The interior of this mineralising body displayed alternating concentric bands of higher and lower BSE intensity together with a nanoparticle-like texture in SE images (well correlated to the texture in BSE mode) resembling Liesegang rings. Such ring-like structures were also located in block-face SEM imaging.

#### 7. Methods

##### Optics sample preparation

For *Elysia viridis,* individuals were sedated by adding isotonic $MgCl_2$ solution dropwise to seawater, and a cover slip was placed on the dorsal surface with spacers on either side to level the surface of the animal. The spectra were taken as quickly as possible to limit exposure of the symbiotic chloroplasts to high light, which could cause damage. Due to the polycrystalline nature of the photonic crystals found in this species, multiple spectra were taken across each calcophore (since the spectral collection area is < 3 µm wide).

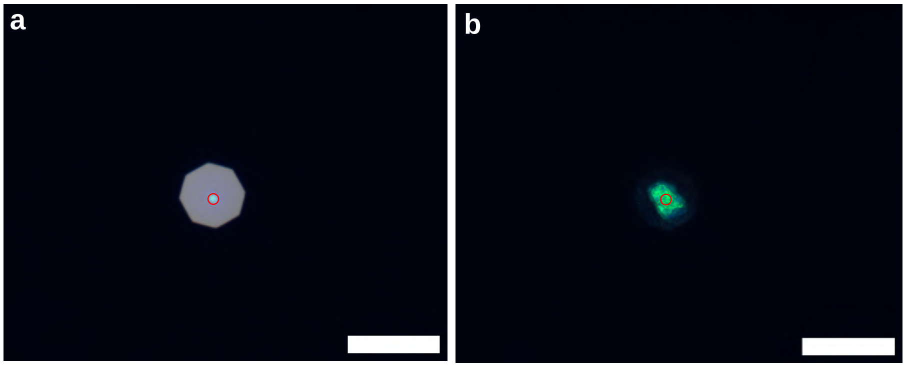


**Supplementary Figure 26**

Area illuminated with a lamp to define the spectral collection area. **a**) Shows the illumination condition, reflection here is from the silver mirror used as a reference. **b**) Shows the collection condition with an example calcophore. Samples are illuminated with a 50 µm optical fibre. The red ring is added on the PixelLink software to know which region of the sample spectra are being collected from. Images taken on Zeiss Axio Scope A1 with a Zeiss W N-Achroplan 63X objective. Scale bar, 50 µm.

Samples were imaged on a Keyence VHX-7100 digital microscope before being transferred for spectra. Seawater was used as the medium between the water immersion objective and the cover slip to ensure index matching. Spectra were taken in the Köhler illumination configuration, with the aperture diaphragm nearly fully closed to reduce stray light from undesired areas. An image of the fibre collection area was taken prior to spectra using an external lamp, and a circle representing this area was overlaid on the PixelLink imaging software to accurately know from which area spectra were being taken (**Supplementary Figure 26**).

##### Optical Microscopy and Micro-Spectroscopy

Micro-spectroscopy was carried out using a customised Zeiss Axio Scope A1 microscope, fitted with a Pixelink PL-D725CU-T high frame rate camera, which was colour calibrated with a white diffuser (Labsphere USRS-99-010). A Zeiss HAL100 halogen lamp was used as a light source. For acquiring spectra, the microscope was coupled to an Avantes AvaSpecHS2048 spectrometer using an Avantes FC-UVIR50 (50 µm core diameter) optical fibre. Spectra were taken using a water-immersion objective lens at 63x, depending on the sample (Zeiss W N-Achroplan 63X (NA 0.9, FWD 2.4 mm). Spectra were referenced to a protected silver mirror (PF10-03-P01, with >97.5% reflectance for 450nm - 2µm).

#### Digital microscopy and photography

A Keyence VHX-7100 digital microscope was used to take high magnification images of live sea slugs using either the full ring or partial ring modes (**Supplementary Figure 27**). Full ring mode is equivalent to dark field, in which the sample is illuminated from a ring of light on the outside of the objective, and light is collected from the centre. This ring can be split into 4 separate illumination angles for 'partial ring' imaging. A Nikon D7500 SLR camera was used to take macroscopic images of the sea slugs.

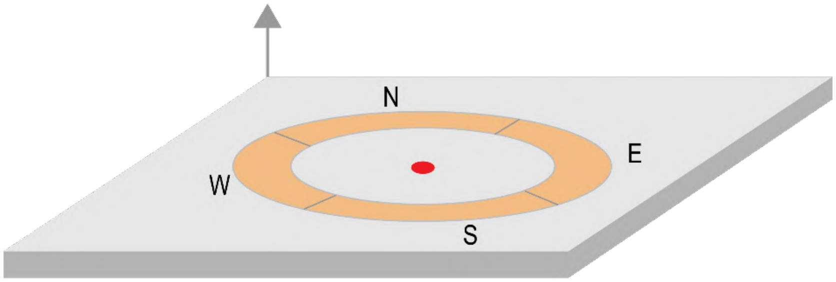


**Supplementary Figure 27**

Schematic showing the illumination configuration of digital microscopy using a Keyence VHX-7100. The sample is illuminated either by the entire ring (full ring mode), or any one of the NESW directions (partial ring mode). Reflected light is collected from the centre (marked by a red circle).

#### K-space imaging

Due to the size of the area where the structural colour in *Elysia viridis* is found (10 - 30 µm diameter), macro-scatterometry techniques such as angular resolved spectroscopy with an optical goniometer can be challenging, so micro-scatterometry (conoscopic imaging) was used to acquire useful information on the angular distribution of the scattering properties of the photonic structures. To have access to this information K-space imagining or conoscopic imaging was performed with a Betrand Lens.

A slider with a focusing Bertrand lens (Zeiss 453671-0000-000) was inserted into the customizable Zeiss Axio Scope A1 microscope above the carousel, such that it does not interfere with the light incident on the sample. Illumination for real-space images was from a Zeiss HAL100 halogen lamp, whilst for k-space images, an Ocean Optics HPX2000 xenon light source was coupled to an Avantes FC-UVIR50 (50 µm core diameter) optical fibre. Prior to aligning the Bertrand lens, the real-space image was focused by removing the slider and focusing on a silver mirror. The Bertrand lens was then inserted, and the back focal plane (Fourier plane) image was focused to align the lens using a Thorlabs transmission grating GT13-12, 1200 Grooves/mm (a = 833 nm). The sample was then inserted, and images were taken switching back and forth between the real and reciprocal space images. The exposure time was increased for reciprocal-space imaging because the scattered signal was dominated by reflections normal to the sample surface. K-space imaging was performed with a Zeiss and W N-Achroplan 63X (NA 0.9, FWD 2.4 mm), firstly because the high magnification allows scattered signal to be isolated from structurally coloured regions, and secondly because the high numerical aperture captures a greater range of scattered angles.

**Optical Principles of K-space imaging**

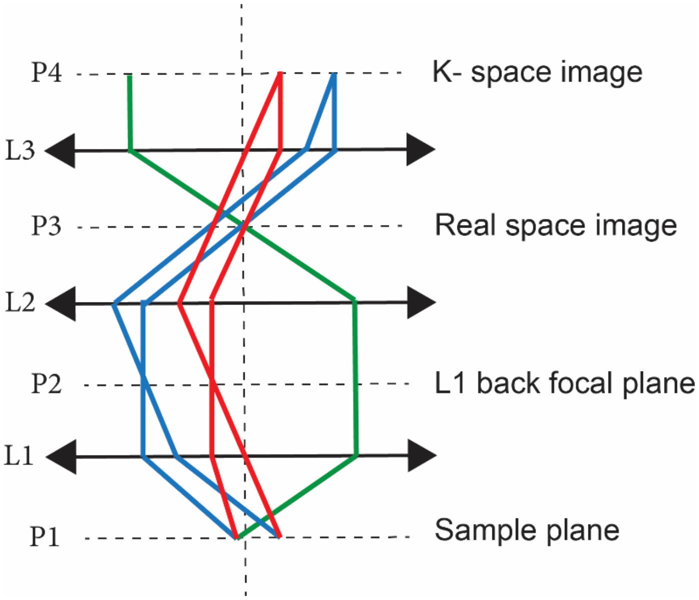


**Supplementary Figure 28**

Schematic showing the principle of K-space imaging. L3 is the inserted Bertrand lens. Light leaving the sample at the same angle, but different positions end up at the same point in the K-space image (2 red, or 2 blue lines). Light leaving the sample at different angles, but the same position ends up at a different position in the K-space image (e.g. green and blue line).

Reproduced with permission from [19]

K-space or conoscopic imaging allows access to information on the directionality of the scattered beam, that is, to image the back focal (Fourier) plane directly, otherwise known as the K-space (or reciprocal space) distribution.

The principle of micro scatterometry using conoscopic imaging in reflection mode is shown in **Supplementary Figure 28** (for simplicity, the illumination paths are not reported in the illustration. The different coloured rays correspond to reflected light leaving the sample at different angles, from different points). The light reflected from the sample is collected by an objective lens L1 (63X, NA = 0.9). The sample is focused on real space such that the focal plane of L1 is P1, this means that the reflected light rays are focused onto the back focal plane of L1, P2. The second lens L2 is positioned in telescopic configuration with respect to L1 such that an image of the sample plane is created at P3. The Bertrand lens (L3) is then mounted in the telescopic configuration with respect to L2, such that an image of the back focal plane P2 is formed at P4. To simplify this diagram all lenses have been displayed with identical focal lengths, and the distances between the lenses are shown to be double the focal length. This diagram demonstrates that two reflected rays leaving the sample at different positions will end up at the same radial position in the K-space, and that the higher the angle of scattering, the larger the radial distance in the K-space image. This is visible in **Supplementary Figure 28**, the two blue (and two red) rays arrive at the same K-space position despite being at different sample positions due to the light leaving the sample plane (P1) at the same angle, whilst light rays leaving the sample at the same position will only arrive to the same K-space position if they leave the sample plane at the same angle. The blue rays leave the sample at a greater angle of reflection and therefore end up further from the centre of the K-space image. This technique therefore allows one to determine the correlation between scattered angle, colour, and intensity. [19]

Different positions in the back focal plane image correspond to different scattering angles from the sample. By projecting the K-space image onto a camera, one can image directly the different Fourier components of the reflected light, in colour, intensity and scattering angle. The farthermost positions in the K-space image correspond to the maximum NA of the objective lens.

**TEM Imaging**

A JEM 2100Plus (JEOL, Japan) transmission electron microscope operated at 200 kV was used for TEM imaging and in 4D STEM imaging. It is equipped with a Gatan RIO16 CMOS camera (Gatan, USA). To enhance the imaging contrast, a 60-µm objective aperture was employed. Low-dose imaging was done in the Low-Dose mode of the SerialEM program (Mastronarde, 2003). TEM and 4D STEM were done with assistance from Yu Ogawa, University of Grenoble.

**TEM Sample preparation**

A stainless steel dewar was filled with liquid nitrogen, a small container (a plastic 2 ml Eppendorf tube) was then partially submerged in the liquid nitrogen and allowed to cool. A brass wire was used to suspend the small container in this position. The container was filled with ethane gas, which was then liquefied.

For *Elysia viridis*, an individual was decapitated, and the head was placed back into an aquarium to regenerate. A small piece of tissue (approximately 1 x 2 mm) was removed from the parapodia of the body of that individual with a scalpel (immediately after decapitation to ensure the sample was as close to its in vivo state as possible) and submerged in 2% glutaraldehyde in cacodylate buffer for 20 minutes. The tissue was then placed, overhanging from the thin end, onto a piece of filter paper which had been cut into a trapezium shape of length ~5 mm, tip width ~2 mm, and base width ~3 mm (**Supplementary Figure 29**). Filter paper was used to partially blot the excess seawater, which would result in more time-consuming trimming later, and over-blotting was avoided by immediately freezing the sample after placing it onto the filter paper. Holding the filter paper with inverted tweezers, the sample was then plunged into the liquid ethane for approximately 10 seconds to ensure that it is fully frozen, after which it was transferred to a polystyrene container with liquid nitrogen until mounted for cryo-ultramicrotomy.

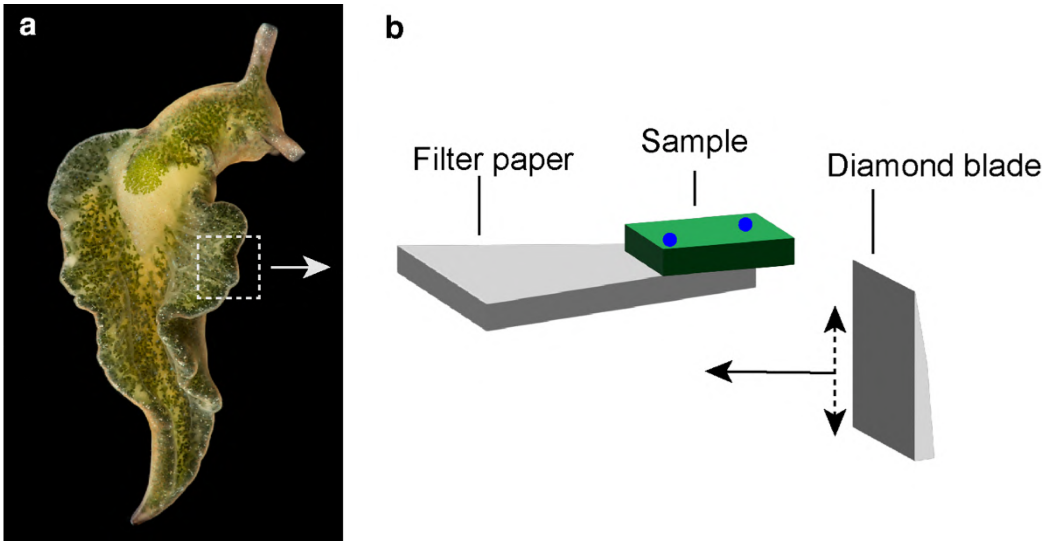


**Supplementary Figure 29**

a) Example image of an *Elysia viridis* individual with a region of interested highlighted, taken with a Nikon D700 and Nikon macro-objective (105 mm).

b) Schematic of TEM sample preparation. The region of interest in is removed with a scalpel and added to a trapezium shaped piece of filter paper before freezing. Microtoming with a diamond blade is done towards the filter paper.

For fixed tissue TEM and SEM imaging, an entire individual was sedated by adding isotonic $MgCl_2$ (3.5%) dropwise to seawater until movement had stopped. The individual was then placed in cacodylate buffer with 2 % glutaraldehyde for 5-6 hours. Prior to embedding, the sample was stained with 1% osmium tetroxide for 2 hours, then washed 4 times in buffer before being dehydrated in an ethanol series. The sample was then infiltrated with ethanol:Epon resin (50:50) overnight, with resin then exchanged every 24 hours for 3 days. To polymerise the resin, a DMP accelerator was added, and the samples were warmed to 60 °C for 24 hours. The following table gives the details of the process:

| **Step** | **Time** |
|---|---|
| 2% Glutaraldehyde | 5-6 hours |
| Buffer wash | 30 min |
| Buffer wash | Overnight |
| Osmium | 2 hours |
| Buffer wash | 5 min |
| Buffer wash | 10 min |
| Buffer wash | 10 min |
| Buffer wash | 30 min |
| 30% ethanol | 10 min |

| | |
|---|---|
| 50% ethanol | 10 min |
| 70% ethanol | 30 min |
| 90% ethanol | 30 min |
| 100% ethanol | 30 min |
| 100% ethanol | 30 min |
| 100% ethanol | 1 hour |
| 50% ethanol, 50% Epon resin | Evaporate open over night |
| 100% resin | 3 hours |
| 100% resin | 3 hours |
| 100% resin | 3 hours |
| 100% resin | 1 day |
| 100% resin | 1 day |
| Resin + DMP | 12 hours |
| Resin + DMP | 12 hours |
| Polymerization | 24 hours |

**Freeze substitution**

The freshly cut tissue was sandwiched individually between two type B gold-coated copper high-pressure freezer carriers (BALTIC preparation, Wetter, Germany) with the addition of seawater. The carriers were positioned in a mirror combination to allow a total cavity thickness of 0.6 mm. The sandwiched samples were cryo-immobilized in an EM ICE high-pressure freezing machine (Leica Microsystems, Vienna, Austria) within 10 minutes after dissection. Freeze substitution was carried out in an automatic freeze substitution unit (Leica Microsystems, AFS). The samples were first transferred under liquid nitrogen to tubes containing freeze-substitution medium composed of 100% ethanol containing 20mM oxalic acid. Tubes were placed in AFS, which was set at −120 °C. They were subsequently warmed to −85 °C at a rate of 17.5 °C/h and held at −85 °C for 103 h, then warmed to −20 °C at a rate of 7.2 °C/h, followed by a hold at −20 °C for 12 h, and a final heating to 4 °C at a rate of 2.7 °C/h, followed by holding at 4 °C for 24 h. The samples were rinsed with ethanol and infiltrated with Embed 812 epoxy resin for 3 days. The resin polymerisation was done for 24 hours at a curing temperature of 65 °C.

**Cryo ultramicrotomy**

The sample was transferred from liquid nitrogen into the ultramicrotome chamber with precooled tweezers (in liquid nitrogen). The non-sample end of the filter paper was then fixed in the clamp of a specimen holder and left until the temperature had equilibrated with the set temperature (-110 °C for the sample and for the knife). An antistatic electrode was placed in the cryochamber and switched on before sample loading. The knife holder was then rotated such that the trimming tool was in the cutting position, which was then moved to the edge of the sample. The sample was then trimmed on either side (in a particular position such that it was trimmed around where a structurally coloured region was, based on observation), to give a flat surface of less than 100 x 100 µm. The knife holder was then rotated to the diamond knife position and brought to the face of the sample. Ultrathin (90 - 100 nm) sections were cut with the diamond knife at a speed of 0.1 mm $s^{-1}$ and transferred to precooled, carbon-coated, and glow-discharged (at 15 mA for 25 seconds with a glow discharger) copper TEM grids using an eyelash. To prevent ice crystal growth after sectioning the grids were transferred to a vacuum chamber where they were brought to room temperature under vacuum. The grids were then ready for TEM imaging. The cryo-chamber used was a Leica microsystems FC7C and the ultramicrotome was a Leica EM UC6. The knives used for sectioning were a trimtool (45°) and a cryo-immuno diamond knife (35°) (both made by Diatome, USA). Cryo ultrathin sections of thickness of about 100 nm were cut and collected on carbon-coated copper grids at -110 °C. The sections were transferred to a vacuum chamber to freeze-dry to minimise the structural damage during thawing.

**TEM**

A low dose (no greater than 4 $e^{-}/\text{Å}^{-1}$) is used for TEM imaging due to the vulnerability of the unfixed, unstained and unembedded biological samples. Imaging was done at room temperature. The sample was mounted in a standard holder and inserted into the TEM, the eucentric height adjusted and the TEM aligned where necessary. Using low dose mode in the SerialEM software, the beam was setup in an area of grid without sample. The low dose mode in SerialEM software (https://bio3d.colourado.edu/SerialEM/hlp/html/about_low_dose.htm) separates the image into 3 areas- ‘Record’, ‘View’ and ‘Focus’. The record area is where micrographs are recorded, and the view area is a low magnification image with the record area at its centre. The focus area is displaced from the record area along its tilt axis and is used to check and correct the defocus without damaging the record area by over exposure during focusing. Images were then acquired at various magnifications. In *Elysia viridis* samples the desired imaging area occupied a small proportion of the area of each section. However, due to the higher electron-absorption density of calcium carbonate, these areas could be observed at relatively low magnification and then zoomed in on for further imaging.

**4D STEM**

Scanning nanobeam electron diffraction (SNBED) images were collected with a JEM 2100F at 200 kV operation voltage, equipped with Cheetah hybrid pixel detector (Amsterdam Scientific Instruments) and a NanoMEGAS ASTAR system. The converged electron nanobeam used as a probe had a pixel diameter of 10 - 25 nm. The detailed description of 4D STEM acquisition conditions can be found in Lim et al. (2023, J. Phys. Chem. Lett).

**STEM EDS**

STEM EDS was done on the same sections as above, but with assistance from Dr Heather Greer in the Department of Chemistry, Cambridge. The TEM used was a Talos F200X G2 (Thermo Scientific) operated at 200 kV. STEM images were collected using a Fischione High Angle Annular Dark Field (HAADF) detector at a camera length of 98 mm. Energy Dispersive X-Ray Spectroscopy (EDS) spectra and maps were acquired using a Super-X EDS detector system, which consists of four windowless Silicon Drift Detector's (SDD).

**SEM EDS**

EDS measurements were conducted on block faces of resin-embedded tissue samples. The block faces were coated with a 10 nm-thick amorphous carbon layer. EDS elemental mapping was performed using an Ultim Max 170 detector (Oxford Instruments, UK) within a Crossbeam 550 FIB/SEM (Carl Zeiss AG, Germany) operated at an accelerating voltage of 7 kV with the specimen at a working distance of 8 mm.

**Cryo Focused Ion Beam-Scanning Electron Microscopy (FIB-SEM)**

The desired areas of tissue were cut from the body of a decapitated *Elysia* viridis body with a scalpel and under a stereo microscope (Zeiss) for accuracy. Immediately after dissection the sample was placed into a B gold-coated copper freezer hat (BALTIC preparation, Wetter, Germany) and 10 wt% dextran (Sigma, 31390) was added as a cryoprotectant. A second freezer hat was added in a mirror configuration to enclose the sample, giving a total cavity thickness of 0.6 mm. Within approximately 5 minutes of decapitation, the sample in this sandwich combination was cryo-immobilised using a high-pressure freezer (HPM100, Leica Microsystems). The frozen sample holder in a sandwich configuration was then mounted into the cryo sample holder of a Leica EM VCM loading station (Leica Microsystems) at -196 °C. A VCT500 shuttle was used to transfer the sample to an ACE600 (Leica Microsystems) for freeze fracture and sputtering. After freeze-fracture, the exposed surface of the sample was sputter-coated with platinum to a thickness of ~8 nm. The VCT500 was then used to transfer the sample to the Zeiss Crossbeam 540 (Zeiss Instruments) for imaging. The VCT500 allowed us to transfer samples at a maximum temperature of -145 °C.

**Serial surface view imaging**

Focused ion beam scanning electron microscopy (FIB-SEM) serial surface view (SSV) imaging was done using a Zeiss Crossbeam 540 (Zeiss instruments). Samples were raised to 5.1 mm (the coincident point of the electron and focused ion beams) and tilted to 54 °. A trench, approximately 120 µm wide and 20 µm long, was milled overnight at 15 nm; Individual 20 µm wide areas were polished with a low ion beam current of 1.5 nA to search for mineralised areas in the back-scattered signal. To cover a larger volume with fewer sections, we only look at every other 20 µm section. After finding such areas, the polishing was completed at 1.5 nA such that the surface was ready for SSV imaging. The electron beam was focused on the polished surface at 2 keV and 50 pA. Depending on the size of the volume being imaged, the x by y pixel size was either set to 8 x 8 nm or 12 x 12 nm. The slice thickness in the z direction was then set such that the voxel size was isometric, the images were then sequentially collected with the 'slice and view' protocol. Stacks were collected with an image in 8-bit greyscale, using the dual channel option. The dual channel option allows for acquisition of images from the mixed InLens /Secondary electron (SE) detector, and a backscattered electron (BSE) detector.

**Image processing and segmentation**

The alignment of stacks, segmentation and image processing were done using Dragonfly 3D World software (Object Research Systems Inc). Both the mixed and BSE signals were aligned automatically with the sum of square differences (SSD) algorithm available in the Dragonfly software; these were then corrected manually to ensure there were no artefacts. The vertical destriping filter was used to remove artefacts due to curtaining (a result of uneven beam milling due to volumetric density fluctuations). Contrast was enhanced with a convolution filter for BSE images, and both a convolution filter and contrast-limited adaptive histogram equalisation (CLAHE) filter for mixed images.

For segmentation of the backscattered images where contrast is high and there are only two regions of interest, a simple Adaptive Gaussian model was trained over 5 images- and used to segment the entire volume. A watershed transform function was used to discretise individual nanoparticles which, in some cases could not be separated due to their proximity. In this case, they were separated manually. This process allows for statistical information about the particles to be obtained. For segmentation of the mixed volumes, a deep learning module in Dragonfly was used with 6 slices for training, and manual readjustment of the regions of interest between each training.

**Mapping**

*Elysia viridis* sea slugs were placed in a small petri dish containing seawater (one animal was mapped at a time). A preprepared solution of $MgCl_2$ (Sigma Aldrich) that was isotonic with seawater (3.5 wt%) was then added drop wise to the petri dish to sedate the slug, the solution was added until the animal stopped moving, the slug was then left for an additional 5 minutes before being transferred to a glass slide. 2 - 5 stacked coverslips (100 µm each) were placed on either side of the specimen, across which another coverslip was placed to level the surface of the animal such that the focal plane would be the same across the surface area, and to ensure that parapodia remain open so that the entire body could be mapped. Images were taken at 170X magnification on a Keyence VHX-7100 digital microscope, using the automatic tiling process built into the software an overall image of the surface of the animal was stitched together to give a high-resolution map. The slide was then flipped over such that is ventral side was facing the objective, and a new tiled image was taken. As soon as imaging was finished the slug was returned to its aquarium where sedation would wear off within 5 minutes and normal activity resumed.

Each image was uploaded to the software QGIS Desktop 3.30.0, and calcophores were manually selected by creating a new point layer for each colour, which was overlayed on top of the microscope image. All layers were given the same co-ordinate reference system to ensure relative distances were accurate (Lisbon, EPSG:4207). Attempts were made to automate this process using a random forest model, however this proved to be inefficient due to the large array of colours, individual calcophores containing multiple colours, proximity of calcophores in clusters, and other features of the slugs containing similar colours. The colours were split into 5 categories: blue, green, orange, red and white. Of course, this is a simplification of the actual colour palette observed in an animal and overlooks the fact that multiple colours can occur within a structurally coloured region due to photonic lattice orientation in a polycrystalline photonic structure. However, for the purposes of this part of the study, 5 colours sufficiently represent the appearance of the animals, and the wavelengths that the calcophores are reflecting. The wavelengths are correlated to the density of nanoparticles inside one calcophore and therefore mapping the colours helps to understand how the calcophores are formed.

After manually assigning a colour to each calcophore as part of a point layer, the total number of each colour was counted automatically in QGIS by the number of points in that layer. Based on these maps, and looking at over 30 animals under the microscope, colour distributions became clear. To quantify these distributions the total surface of the slug (dorsal and ventral) was split into 6 areas, that are on the dorsal side: the heart, the kidney, the bulk and the edge, and on the ventral side: the centre, the bulk and the edge (the edges were combined to give one area). For each slug these areas were defined on QGIS by drawing a polygonal shapefile layer manually, automating this process was not possible because of the compliant nature of the slugs' bodies, morphology varied a lot each time and as such the most accurate way to define these areas was by eye. The number of each colour was counted within each of the specified areas. When separated by colour and area in this way some clear patterns emerge.

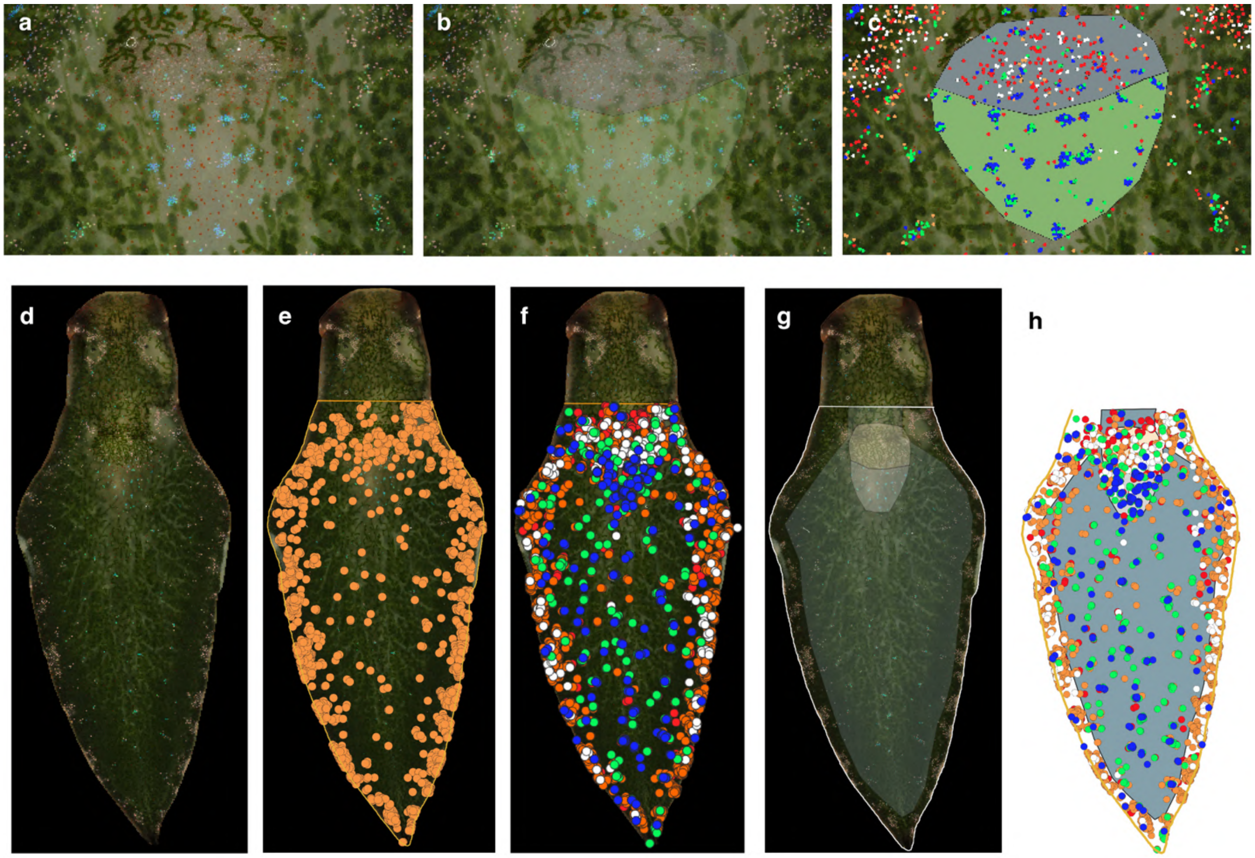

**Supplementary Figure 30**

The process of mapping calcophores across the body of *Elysia viridis* using QGIS software. Points are added manually to a point layer on top of the microscope image for each colour. The surface is then partitioned into 6 areas for each individual and using the QGIS software the number of each coloured point is counted within each of these areas.

a-c) Mapping process for pericardial region.

d-h) Mapping process for entire dorsal surface.

Images of 9 individuals were taken on a Keyence VHX-7100 digital microscope at various magnifications, without sedation, to map the position and number of calcophores around the animal's eye. The calcophores were manually assigned positions by creating a new point layer- colour was not considered, as every calcophore in this region was red/orange/white. To correlate the position and number of these calcophores to the eye itself, the ‘Shortest line between features’ algorithm in QGIS was used. This algorithm creates a new line layer consisting of multiple lines connecting the shortest distance from hubs in a point layer (calcophores) to another layer (the centre of the eye). The output line layer can then be exported as a distance matrix, which can be normalised to a scalebar to give actual distances. It should be noted that even if the surface of the animal is not flat, (and this distance algorithm uses Cartesian co-ordinates, which don’t account for curved surfaces), the error is considered minor. The same algorithm was used to measure the relative distances of different coloured calcophores along a vessel from the kidney to the edge of the animal.

The degree of clustering of calcophores was measured using the nearest neighbour index algorithm in QGIS. Which, for a given area, takes the ratio of the mean distance between calcophores in that area, and the expected mean distance between calcophores if they were randomly distributed in that area.

In order to see whether calcium concentration in seawater has an effect on the total number of calcophores, or the ratio/positions of the different coloured calcophores a solution of 2X calcium concentration (800 ppm) seawater was made up by using $CaCl_2$ solution (calcium chloride was the salt of choice, as seawater is already a highly chlorinated medium due to NaCl, so the addition of more has a minor impact on total concentration). Regrowth of an animal was used as a proxy for growth from a larva (since we were unable to breed the animals). The head of the decapitated individual was placed in the 2X Ca sweater solution with algae and regrew a new body. After 87 days the individual was imaged, and calcophores were counted. This was then compared to an individual which had been decapitated and regrown in normal seawater. Both slugs were imaged before regrowth for a fair comparison, showing a change in the proportion of each colour rather than an absolute change, which is more susceptible to differences between individuals. The body regeneration of a decapitated slug was monitored by regular imaging and measuring the increase in body length over time over the course of 2 months. The initial formation of calcophores was monitored by regular imaging (daily) at magnifications of up to 200X.

**Fluorescence microscopy**

Fluorescence imaging of live individuals was undertaken following protocols from the literature for Zebrafish larvae. Individuals were sedated by adding magnesium chloride solution dropwise before transferring to a 0.2 % Calcein (Cayman chemical, 1461-15-0) in filtered (0.2 µm) seawater solution for 10 minutes. The animals were then rinsed 3 times in seawater for 5 minutes each and placed on a slide, under a cover slip for imaging. Confocal imaging was done with both ZEISS LSM 880 (whole individual, using tiling) and ZEISS LSM 980 with Airyscan 2 (high magnification).

For immunofluorescence imaging of fixed tissue, individuals were sedated and decapitated (the head regenerated a new body) before dissecting to the desired volumes. These volumes were then fixed for 4 hours in a 4% solution of PFA (paraformaldehyde, Sigma Aldrich) in PBS (phosphate buffered saline) with added NaCl to make a solution isotonic with seawater. The samples were rinsed 3 times in PBS and left overnight in a 30% sucrose solution (as a cryoprotectant) in PBS until the tissue sunk. The samples were then embedded in Neg 50™ (Fisher Scientific) and plunged into liquid nitrogen before storage at -80 $^{o}$C. Samples were sectioned at -20 $^{o}$C in a Leica CM3050 S cryostat to 12-16 µm thick sections and mounted onto Epredia™ SuperFrost Plus™ microscope slides. The mounted sections were washed 3 times for 5 minutes in PBS and then incubated in 0.2% Calcein in PBS for 20 minutes, followed by washing 3 times again in PBS. The sections were then incubated in a 1X working solution of Phalloidin-iFluor 594 Reagent (abcam) in PBS for 40 minutes and subsequently washed 3 times in PBS for 5 minutes. The sections were then incubated in a solution of NucBlue™ Fixed Cell ReadyProbes™ Reagent (DAPI, Fisher Scientific) in PBS (2 drops per ml) for 20 minutes and washed again with PBS 3 times. The stained sections were placed under a cover slip and immediately imaged. Imaging was done on both an Invitrogen™ EVOS™ M5000 Imaging System (with assistance from Edward Wills in the Ladds group, University of Cambridge) and a Leica SP8 Confocal Microscope. Imaging was done with the following excitation laser wavelengths and detector wavelengths. DAPI: Ex (405 nm), Det (415 - 470 nm). Calcein: Ex (488 nm), Det (500 - 540 nm). Phalloidin: Ex (552 nm), Det (610 - 630 nm).

#### Holotomography

Samples of *Elysia viridis* for holotomography imaging were prepared using dissected tissue (approximately 0.5 x 0.5 mm) immediately after decapitation of a sedated individual. The samples were compressed under a cover slip to ensure they were thin enough for accurate refractive index measurements and placed in a Tomodish sample holder for imaging. Samples were then imaged using a Tomocube HT-X1™. The principle of 3D refractive index measurement using holotomography is that a rotating incident illumination is split into two, one of the beams goes through the sample and the two then recombine. The phase difference between these two beams depends on the passage of the beam through the sample which depends on the refractive index of the material in the sample. The holotomography images were analysed using the TomoAnalysis software.

#### Raman spectroscopy

Raman spectroscopy was performed on cryo-cut, unfixed, unstained 14 µm thick sections of sea slug tissue. Samples were cut at -20 ° using a Leica CM3050 S cryostat. An alpha300 R - Raman Imaging Microscope (Oxford Instruments) equipped with a UHTS spectrometer was used to take Raman spectra. Spectra were taken with a 100X objective (Nikon) in confocal mode. The desired region of interest was located on the sections using a Keyence microscope in full ring mode. The sample was then transferred to the Raman microscope, and the desired region of interest was measured in at least 3 positions with an integration time of 60 seconds for each. These positions were averaged and processed using the Project *FIVE*+ software (Oxford Instruments). For Raman mapping an integration time of 5s was used due to time constraints. Raman spectroscopy was done with assistance from Dr Tobias Priemel.

#### Sample collection and ethics

*Elysia viridis* specimens were collected at low tide in Le Croisic, France and Goodwick, Wales, using either a small fishing net or a paintbrush to remove the animals from algae. Algae were also collected as food for the animals from their respective habitats. Individuals from Wales were kept in aquaria in Cambridge, UK, whilst those from France were kept in aquaria in Golm, Germany. Lloyd Nelmes (Sea Trust) and Bruno Jesus (University of Nantes) assisted in sample collection. The aquaria were filled with a mixture of seawater from the habitats of the animals and artificial seawater. Animals were kept at 18 °. All specimens were handled in accordance with best practices, aiming to reduce animal suffering where possible. This includes sedating of animals prior to termination or decapitation, termination by flash freezing where possible and regular replacement and aeration of seawater.

#### Double-ended probe

A double-ended reflection probe (Ocean Optics R200-7-SR) was used to measure reflection from *Elysia viridis* in the open and closed parapodial positions. This experiment was done on an individual who had already died in order to prevent stress to a live animal. The body was positioned using glass slides, and the probe was positioned close to the animal to give an approximately circular illumination area of 200 µm diameter. Spectra were normalised to a silver mirror.

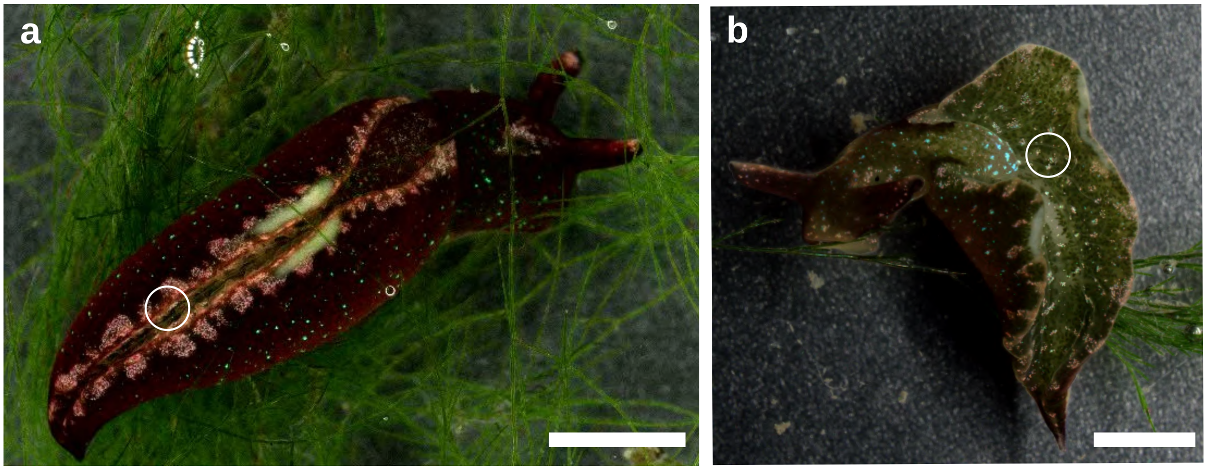


**Supplementary Figure 31**

The closed (**a**) and open (**b**) parapodial positions of *E. viridis* triggered by high and low light conditions. Circles indicate the positions of double ended probe light illumination and collection. Scale bars, 3 mm, 2mm.

##### Optical simulations

The 5 x 5 x 5 µm correlated and polycrystalline particle assemblies were generated using the PolydisperseMD v.2.9 plugin[20] for HOOMD (v.2.9.4)[21] using Langevin integrator (kT = 0.05) and polydisperse 12-0 softcore pair-potential, and the expanded FCC lattices with Matlab. The particle coordinates and diameters were then imported to Ansys Lumerical finite-difference time-domain (FDTD) simulator and depending on the case either plane wave or Gaussian beam was used as the illumination source and reflection monitor was placed above source. A perfectly matched layer (absorbing) boundary condition were used for the simulation box boundaries, and the measured intensities were integrated over the monitor area.
